# 2-Cys peroxiredoxins facilitate phosphatidylglycerol unsaturation and contribute to PSII recovery from high-light photoinhibition

**DOI:** 10.64898/2026.11.16.766973

**Authors:** M. Luisa Hernández, Julia Jiménez-López, Fernando Rodríguez-Marín, Valle Ojeda, Alicia Gálvez-Ramírez, Francisco J. Cejudo, Juan M. Pérez-Ruiz

## Abstract

The level of fatty acid unsaturation of chloroplast lipids is a key determinant of plant acclimation to different environmental stresses. Though the enzymes catalysing fatty acid desaturation are well known, the regulatory mechanisms linking their activity to environmental signals remain poorly understood. In this work, we have addressed the role of chloroplast 2-Cys peroxiredoxins (Prxs), thiol peroxidases that play a central role in redox regulation, in ω-3 fatty acid desaturation in lipids of the organelle, and their contribution to plant acclimation to environmental stress. Genetic and lipidomic analyses of Arabidopsis (*Arabidopsis thaliana*) mutants lacking 2-Cys Prxs and/or the chloroplast ω-3 fatty acid desaturases FAD7 and FAD8 revealed that 2-Cys Prxs are required for the efficient synthesis of trienoic fatty acids in thylakoid membrane lipids, particularly in phosphatidylglycerol (PG). Different approaches show the *in vivo* interaction of 2-Cys Prxs with FAD7 and FAD8. However, no redox changes of these desaturases were observed regardless the contents of 2-Cys Prxs, thus indicating a regulatory mechanism independent of the classical cysteine-based redox switch. The combined absence of 2-Cys Prxs and chloroplast ω-3 desaturases severely compromised recovery from high-light photoinhibition, indicating that both systems act concertedly to maintain photosystem II (PSII) function under stress. Together, our results uncover a novel role of 2-Cys Prxs controlling the level of chloroplast PG *cis*-unsaturation and raise the possibility that the chaperone/scaffold activity of these enzymes links membrane lipid remodelling to photosynthetic stress response.

## Introduction

Lipids, as essential structural and functional components of cellular membranes, play critical roles in plant development and adaptation to variable environmental conditions. A key aspect of plant adaptation to the environment is the modulation of membrane fluidity, which is highly dependent on the composition of the membrane lipid bilayer, remarkably the degree of unsaturation in their fatty acid acyl groups (Li et al., 2016; Li et al., 2020; Cook et al., 2021; Henschel et al., 2024; Shomo et al., 2024). In plants, the most abundant class of membranes lipids are glycerolipids; however, important differences in lipid composition characterize the different plant cell compartments, while extraplastidial membranes are enriched in phospholipids, chloroplast membranes have glycolipids as main constituents (Hölzl and Dörmann, 2019). Among chloroplast glycolipids, the uncharged galactolipids monogalactosyldiacylglycerol (MGDG) and digalactosyldiacylglycerol (DGDG) are the main components of thylakoid membranes, hence providing the bulk of the lipid matrix where photosynthetic complexes are embedded. Lower contents are found of the anionic lipids sufoquinovosyldiacylglycerol (SQDG) and phosphatidylglycerol (PG), this one being the only phospholipid component of thylakoid membranes (Hölzl and Dörmann, 2019). An additional feature of chloroplast lipids is the high proportion of trienoic fatty acids (TAs). α-linolenic (18:3) and hexadecatrienoic (16:3) acids typically comprise about 70% of total fatty acids, both constituting more than 90% of the MGDG fatty acids (Hölzl and Dörmann, 2019). This unique membrane system not only serves as a structural scaffold for the chloroplast photosynthetic machinery, hence for photosynthetic performance, it is also relevant for the metabolic and signalling networks involved in chloroplast biogenesis, and the acclimation of these organelles to different environmental stresses (Kobayashi, 2016; Hölzl and Dörmann, 2019; Hernández and Cejudo, 2021; Fujii et al., 2025).

In higher plants, *de novo* fatty acid biosynthesis is initiated in the chloroplast, and glycerolipid assembly can be accomplished by two different pathways. PG is mostly synthesised by a metabolic pathway entirely located in the chloroplast, the so-called prokaryotic pathway (Li-Beisson et al., 2013; Hurlock et al., 2018). The galactolipids synthesised by this pathway are characterized by the presence of C16:3 fatty acid in the *sn*-2 position of the glycerol backbone. An alternative metabolic pathway for chloroplast lipids synthesis, the so-called eukaryotic pathway, uses glycerolipids assembled in the endoplasmic reticulum (ER) that return to the plastid and serve as substrates for galactolipid synthesis into the organelle. The ER located acyltransferases have a high specificity for C18 fatty acids, thus the presence of C18:3 fatty acids in the *sn*-2 position of glycerol has been traditionally used a signature for eukaryotic galactolipids. However, evidence from *Chlamydomonas* and Arabidopsis shows that ER-derived lipid precursors can acquire plastid-like acyl compositions and undergo extensive remodelling (Mugume et al., 2026). Additionally, the relative contribution of the prokaryotic and eukaryotic pathways varies among species. In some plants, like Arabidopsis, both pathways contribute almost equally to galactolipid biosynthesis, while in other species, such as soybean or wheat, galactolipids are exclusively synthesised by the eukaryotic pathway and, thus, lack C16:3 fatty acids (Li-Beisson et al., 2013).

Regardless of the biosynthetic pathway, fatty acids esterified to membrane lipids undergo further desaturation by membrane-bound fatty acid desaturases (FADs), which vary in cellular location, substrate specificity, and electron donor systems (Shanklin and Cahoon, 1998). Microsomal ω-6 (FAD2) and ω-3 (FAD3) desaturases are localized in the ER, use phospholipid as substrates and NADH and cytochrome *b_5_*, via an NADH-cytochrome *b_5_* reductase, as electron donors. In contrast, the chloroplast ω-6 (FAD6) and ω-3 (FAD7/8) use primarily glycolipids as substrates and ferredoxin and NADPH, via a ferredoxin-NADP reductase, as electron donors. In Arabidopsis, a single nuclear gene encodes for each chloroplast ω-3 FAD, FAD7 (Yadav et al., 1993) and FAD8 (Gibson et al., 1994), the enzymes responsible for the desaturation of dienoic fatty acids (DAs) to TAs in chloroplast lipids. Despite their high homology and the fact that both enzymes have similar activity in the same subcellular compartment, the functions of the two chloroplast ω-3 FADs are not redundant under standard growth conditions. While FAD7 is highly abundant in leaves and primarily responsible for the desaturation of hexadecatrienoic (16:2) and linoleic (18:2) acids in galactolipids, FAD8 is a low abundant enzyme that exhibits higher affinity for the PG-18:2 substrate (Browse et al., 1986; McConn et al., 1994; Román et al., 2015).

The non-redundant functions of both chloroplast ω-3 FADs in the biosynthesis of TAs in chloroplast lipids could be extended to their contribution to plants response to environmental stresses. In support of this notion, a highly coordinated response of both enzymes to stress signals has been reported in Arabidopsis, FAD7 showing higher specialization in the response to biotic stress, and FAD8 in the response to temperature changes (Matsuda et al., 2005; Román et al., 2015; Soria-Garci et al., 2019). Although transcriptional regulation of genes encoding chloroplast ω-3 FADs seems crucial for the response to biotic and abiotic stress (Soria-Garci et al., 2019), a post-translational modification has also been reported to destabilize FAD8 in response to high temperature, the decreased content of the enzyme leading to a decrease in TAs levels (Matsuda et al., 2005). Though the relevance of these enzymes in plant acclimation is well recognized, the current understanding of the molecular mechanisms that allow chloroplast ω-3 FADs to modulate thylakoid membrane TAs levels in response to environmental stresses remains limited.

To address these issues, our group is focussing on chloroplast redox regulatory systems. Chloroplasts harbour a complex redox system in which the NADPH-dependent thioredoxin reductase (NTRC) controls the redox balance of 2-Cys peroxiredoxins (Prxs), thiol-dependent peroxidases that maintain the reductive capacity of thioredoxins, thereby, the activity of redox-regulated enzymes (Pérez-Ruiz et al., 2017; Cejudo et al., 2021). Therefore, 2-Cys Prxs play a central role in the interplay of reducing and oxidizing signals that adjust the redox state of the organelle to changes in light intensity and darkness (Pérez-Ruiz and Cejudo, 2026). Accordingly, the Arabidopsis *2cpab* double mutant, which lacks the two 2-Cys Prxs, A and B, present in the chloroplast, shows delayed growth and sensitivity to high light intensity (Awad et al., 2015). Interestingly, this mutant also shows defective chloroplast biogenesis during embryo (Gallardo-Martinez et al., 2023) and cotyledon development (Ojeda et al., 2021), this phenotype being highly aggravated by high temperature (Rodriguez-Marin et al., 2026). Moreover, the recent finding that 2-Cys Prxs contribute to chloroplast ω-3 FADs functions (Hernández et al., 2024), links the chloroplast redox status to the TAs levels in thylakoid membrane lipids, thus suggesting their function in chloroplast lipids remodelling in response to environmental stresses. Our previous results demonstrated the presence of 2-Cys Prxs in large chloroplast membrane protein complexes containing FAD8. The absence of 2-Cys Prxs compromises FAD8 stability, the lower content of this enzyme causing decreased levels of 18:3 in PGs in the *2cpab* mutant. However, we could not rule out the possibility that FAD7 function was also influenced, since TAs levels in the main thylakoid lipids were lower in the *2cpab* mutant than in the *fad8* mutant, even though the transcript levels of *FAD7* genes were unaffected (Hernández et al., 2024). Altogether, these results suggest a relevant function of 2-Cys Prxs in the control of the level of fatty acid unsaturation in plant chloroplasts, which might be essential for plant development and acclimation to environmental stresses. In this work we have addressed this issue by analysing the functional relationship of 2-Cys Prxs with FAD7 and FAD8. To that end, multiple mutants of Arabidopsis combining the deficiencies of 2-Cys Prxs, FAD7 and/or FAD8 were generated. Based on these results, we identify a key function of 2-Cys Prxs in the modulation of the level of unsaturation of PG fatty acids, exerted via the concerted action on FAD7 and FAD8. Interestingly, this function of 2-Cys Prxs is more relevant for plant response to high light intensity than acclimation to long-term cold treatments.

## Materials and Methods

### Plant material and growth conditions

The ecotype Columbia-0 (Col-0) of *Arabidopsis thaliana* was used as wild type (WT) and to generate the different mutant and transgenic lines used is this study, as described in Supplemental Tables S1 and S2, respectively. T-DNA insertion mutants *2cpa* (Kirchsteiger et al., 2009), *2cpab* (Ojeda et al., 2018), *fad7* and *fad8* (Román et al., 2015) were previously reported. The double mutant *fad7-fad8* (McConn et al., 1994) was derived by manual crossing from single ethyl methane sulfonate (EMS) mutants, with contained point mutations G422A and G446A resulting in W for STOP codon substitutions in residues 141 and 149 of the AtFAD7 and AtFAD8 proteins, respectively (Román et al., 2015). The *2cpa* and *fad8* single mutant were manually crossed to obtain the double *2cpa-fad8* mutant. Mutants *fad7*, *2cpa-fad8* and *fad7-fad8* were subjected to gene base editing using CRISPR/Cas9 technology with RNA guides (gRNA) designed to target the *2-Cys Prx A* (gRNA 2CPA.ex1 and gRNA 2CPA.ex2) and *2-Cys Prx B* (gRNA 2CPB.ex6) genes (Fig. S1A). The specific gRNAs were designed using the web tool CRISPOR (Concordet and Haeussler, 2018) and cloned into pHEE401-E vector, which has an egg-cell specific promoter (Wang et al., 2015). The different mutants lines transformations were carried out via *Agrobacterium tumefaciens* GV3101 using the floral dip method (Clough and Bent, 1998). The selection of transgenic lines was performed on plates containing Murashige and Skoog (MS) medium containing 0.35% (w/v) Gelrite (Duchefa) and 25 µg mL^−1^ hygromycin. The primers used for genotyping T-DNA insertion mutants and for the amplification and sequencing of gRNA complementary sequences are listed in Supplemental Table S3. Edited lines, in which the target genes (*2-Cys Prx A* and/or *2-Cys Prx B*) were mutated, were propagated and selected for hygromycin sensitivity to obtain stable homozygous mutant lines. Sequence analysis allowed the identification of two independent lines containing mutations in the *2-Cys Prx A* and *2-Cys Prx B* genes in the *fad7*, *2cpa-fad8* and *fad7-fad8* genetic backgrounds. The *2cpab-fad7*#1 and *#2* lines, with T-DNA insertion in the *FAD7* gene (Fig. S2A, B) displayed a one nucleotide insertion and six nucleotides deletion in the *2-Cys Prx A* gene, respectively, and one nucleotide insertion in the *2-Cys Prx B* genes (Fig. S1B). On the other hand, the *2cpab-fad8*#1 and #2 lines, with a T-DNA insertion in the *FAD8* (Fig. S2A, B) and *2-Cys Prx A* genes, displayed deletions of one and five nucleotides, in the *2-Cys Prx B* gene, respectively (Fig. S1B). The quadruple *2cpab-fad7-fad8*#1 mutant, with EMS-induced mutations in the *FAD7* and *FAD8* genes (Fig. S2C), displayed a single nucleotide insertion in the *2-Cys Prx A* gene, and a one-nucleotide deletion in the*2-Cys Prx B* gene (Fig. S1B). Additionally, to obtain the *2cpab-fad7-fad8*#2 line, *2cpab-fad7*#1 and *2cpab-fad8*#1 mutants were manually crossed. Therefore, the quadruple mutant contained T-DNA insertions in the *FAD7* and *FAD8* genes (Fig. S2B), one-nucleotide insertion in the *2-Cys Prx A* gene and one-nucleotide deletion in the *2-Cys Prx B* gene (Fig. S1B).

Transgenic lines WT/FAD8-OE and *2cpab*/FAD8-OE were previously described (Hernández et al., 2024). For the generation of WT/FAD7-OE and *2cpab*/FAD7-OE transgenic lines, a genomic fragment containing the full *FAD7* gene (At3g11170) excluding the stop codon (2,166 bp) was amplified, cloned into the pDONR221, subcloned into the binary vector pEARLYGATE102, which adds CFP and HA C-terminal tags, and transformed into the WT and *2cpab* genetic backgrounds as previously described (Hernández et al., 2024).

Plants were routinely grown on soil in growth chambers under long-day (LD, 16 h light/8 h darkness) or short-day (SD, 8 h light/16 h darkness) photoperiods at 22 °C and 20 °C during light and dark periods, respectively, and a light intensity of 125 µE m^−2^ s^−1^. For cold treatments, plants that were grown during 28 d under SD photoperiod, were moved to a similar chamber with the same photoperiod and light intensity except that temperature was set at 10 °C and 5 °C during light and dark periods, respectively, for a period of 30 additional days. The high light treatment was performed on 3-weeks-old plants grown under LD conditions (125 µE m^−2^ s^−1^) that were exposed to a light intensity of 900 µE m^−2^ s^−1^ at 15 °C for 24 hours.

### Sequence analysis and protein predicted structure analysis

Samples were sequenced by STAB Vida (Portugal). Sequences alignments and analyses were performed using the Benchling platform (https://www.benchling.com).

The amino acid sequences of 2-Cys Prx A (Q96291), FAD7 (P46310) and FAD8 (P48622), excluding their transit peptides according to the annotations in the UniProt database (2023), were used for protein complex structure prediction. Protein complexes were modelled using AlphaFold 3 (Abramson et al., 2024) through the AlphaFold Server (https://alphafoldserver.com), using the default settings. For each prediction, ten copies of the 2-Cys Prx A amino acid sequence and one copy of either FAD7 or FAD8 were provided as independent protein chains to model the potential association of each desaturase with the 2-Cys Prx A decamer. Three independent prediction runs were performed for each protein combination (2-Cys Prx A–FAD7 and 2-Cys Prx A–FAD8), generating five structural models per run and a total of 15 models for each combination. Model confidence was assessed using the ranking score, predicted template modeling score (pTM) and interface predicted template modelling score (ipTM) provided by AlphaFold 3. The predicted structures were visualized and inspected using UCSF ChimeraX (Meng et al., 2023) to examine the spatial arrangement of FAD7/FAD8 relative to the 2-Cys Prx A assembly and identify potential regions of interaction. Intermolecular hydrogen bonds and salt bridges between each desaturase and the 2-Cys Prx A subunits were identified using the *hbonds* command in UCSF ChimeraX. The resulting interactions were examined in the representative models and are summarized in Supplemental Tables S4 and S5.

### Protein extraction, alkylation assays and Western blot analysis

Plant tissues, frozen in liquid nitrogen, were ground to a fine powder using mortar and pestle. For total protein analysis, extraction buffer [50 mM Tris-HCl, pH 7.8, 2.5% (v/v) glycerol, 4 M urea and 2% (w/v) SDS] was immediately added, mixed on a vortex and incubated for 20 min at room temperature (RT) and 5 min at 70 °C. After centrifugation at 16,200 *g* for 5 min, the supernatant was collected, and proteins were subjected to SDS-PAGE under reducing conditions using acrylamide gel concentration of 12%. Resolved proteins were transferred to nitrocellulose membranes and probed with the indicated antibodies. Specific antibodies against NTRC (Serrato et al., 2004) and 2-Cys Prx (Pérez-Ruiz et al., 2006) were previously raised in our laboratory. The HA, CHL27 and CF_1_-γ antibodies were purchased from Agrisera (Sweden). Thiol labelling assays by the alkylating agent methylmaleimide polyethylene glycol (MM(PEG)_24_) were performed as previously described (Pérez-Ruiz et al., 2017).

### Immunoprecipitation

Rosette leaves from Arabidopsis lines WT/FAD7-OE and WT, used as a negative control, were fixed with 1% (v/v) formaldehyde by vacuum infiltration for 35 min. Cross-linking reaction was stopped by adding 300 mM glycine followed by three washes with Milli-Q water. Separation of membrane protein fractions and immunoprecipitation assays were performed as previously described (Hernández et al., 2024).

### Split-Ubiquitin-based yeast two hybrid assay

The pNXgate32-3HA and pMetYCgate vectors were used for the protein-protein interaction studies (Obrdlik et al., 2004). The coding sequences of the Arabidopsis 2-Cys PRX A, 2-Cys PRX B, FAD8 and FAD7 proteins were amplified without their respective stop codons using oligonucleotides listed in Supplemental Table S3. To generate the constructs, pNXgate32-3HA (NubG) and pMetYCgate (Cub) were digested with EcoRI/SmaI and PstI/HindIII, respectively. The linearized vectors were co-transformed with the corresponding PCR products, allowing cloning directly in the yeast strain THY.AP4 (*MATa ura3 leu2 lexA::lacZ::trp1 lexA::HIS3 654 lexA::ADE2*) (Fusco et al., 1999).Transformants carrying pNXgate32-3HA and pMetYCgate constructs were selected on YNB medium lacking tryptophan and leucine, respectively. Plasmids were subsequently transformed into *E. coli*, purified, and sequenced to confirm the correct insertion of the different cDNAs. For the interaction experiments, plasmids were co-transformed into THY.AP4 cells and double transformants were selected on YNB medium lacking leucine and tryptophan (-LW). Then, overnight cultures were prepared in liquid YNB-LW medium and adjusted for serial dilution assays. Ten-fold serial dilutions were spotted onto YNB plates lacking leucine and tryptophan (-LW) or leucine, tryptophan, adenine and histidine and supplemented with 150 µM methionine (-LWAH+M). Plates were incubated at 30 °C for 4 d. KAT1 cloned into the corresponding vectors was included as a positive control.

### Lipid analysis

Total lipids were extracted from leaves of the different Arabidopsis lines using the chloroform/methanol method (Li-Beisson et al., 2013), and lipids separation was carried out by thin-layer chromatography as previously described (Hernández et al., 2008). Fatty acid methyl esters of total lipids or individual lipid classes were produced by acid-catalyzed transmethylation (Garcés and Mancha, 1993) and analyzed by gas chromatography as previously described (Hernández et al., 2024).

### RNA extraction and RT-qPCR

Total RNA was extracted from rosette leaves using the Trizol reagent (Invitrogen) and cDNA synthesis was performed with 1 μg of total RNA using the Maxima first-strand cDNA synthesis kit (Thermo Scientific) according to manufacturer’s instructions. Real-time quantitative PCR (RT-qPCR) was performed using cDNA synthesized as stated above in a IQ5 real-time PCR detection system (Bio-Rad) as previously reported (Pérez-Ruiz et al., 2017). Oligonucleotides used for RT-qPCR analyses are listed in Supplementary Table S3. Expression levels were normalized using *ACTIN2* and *UBQ10* as reference genes.

## Statistical analysis

Statistical analysis (two-way ANOVA and one-way ANOVA) were performed using GraphPad Prism version 11.0.2 (GraphPad Software, Inc.). Principal Component Analysis was carried out using the prcomp function from the R package stats of the R Core Team.

## Results

### The contribution of 2-Cys Prxs to FAD7- and FAD8-dependent fatty acid desaturation

The finding that the lack of 2-Cys Prxs causes altered levels of fatty acid desaturation due to FAD8 instability (Hernández et al., 2024) suggested a possible function of 2-Cys Prxs linking the redox state of the chloroplast to the level of fatty acid unsaturation in lipids of the organelle. It is known that FAD7 and FAD8 activities are not redundant under standard growth conditions, however, the loss or altered activity of one of these enzymes can be compensated for, at least partially, by the other (Román et al., 2015). Thus, the question arising is whether 2-Cys Prxs act concertedly on FAD7 and FAD8 to control the level of thylakoid lipid fatty acid unsaturation. To address this issue, we have generated multiple mutant lines of Arabidopsis combining the absence of 2-Cys Prxs with that of FAD7 and/or FAD8 (Table S1). Due to the genetic linkage between *2-Cys Prx A* (At3g11630) and *FAD7* (At3g11170) loci, as well as between *2-Cys Prx B* (At5g06290) and *FAD8* (At5g05580) loci, obtaining these mutant combinations by crossing available T-DNA insertion lines was not feasible. To overcome this impediment, we performed CRISPR/Cas9 gene editing using gRNAs specifically designed to knockout *2-Cys Prx A* (gRNA-2CPA-ex1 and gRNA-2CPA-ex2) and *2-Cys Prx B* (gRNA-2CPB-ex6) genes (Fig. S1A) in the *fad7* (T-DNA mutant), *2cpa*-*fad8* (double T-DNA mutant), and *fad7-fad8* (EMS generated mutant) genetic backgrounds (Table S1). Sequence analyses allowed the identification of independent mutants *2cpab*-*fad7* (lines #1 and #2), *2cpab*-*fad8* (lines #1 and #2) *and 2cpab*-*fad7-fad8* (line #1), displaying a variety of editing events in the *2-Cys Prx A* and the *2-Cys Prx B* genes, including single- and multiple-nucleotide indels causing frameshift mutations (Fig. S1B). A second quadruple mutant, *2cpab-fad7-fad8* (line#2), was generated by manually crossing the *2cpab-fad7*#1 and *2cpab-fad8*#1 mutants. Immunoblotting analysis confirmed the absence of 2-Cys Prxs in the *2cpab* genetic backgrounds (Fig. S1C) whereas PCR-based genotyping (Fig. S2A, B) confirmed that mutants were homozygous for the T-DNA insertions in the *FAD7* (*fad7, 2cpab*-*fad7* and *2cpab-fad7-fad8*#2 mutants) and the *FAD8* (*fad8* and *2cpab*-*fad8* and *2cpab-fad7-fad8*#2 mutants) genes. Finally, gene sequencing (Fig. S2C) confirmed the EMS-induced mutations in the *fad7-fad8* and *2cpab*-*fad7-fad8#*1 mutants.

When grown under LD photoperiod, the deficiencies of FAD7 and/or FAD8 in the *2cpab* mutant background resulted in growth rates slightly lower than those of the *2cpab* and the *fad7-fad8* mutants, as shown by the fresh weight of rosette leaves (Fig. 1A-B). Accordingly, the photosynthetic performance, as reflected by the maximum photosystem II (PSII) quantum yield (F*_v_*/F*_m_*), which was lower in the *2cpab* than in the WT, showed similar levels in the *2cpab, fad7-fad8* and *2cpab-fad* multiple mutants (Fig. 1C). Overall, these results indicate that the absence of chloroplast ω-3 FAD function has no additional effects on growth and photosynthetic performance than the lack of 2-Cys Prxs, at least under standard growth conditions.

**Figure 1.**
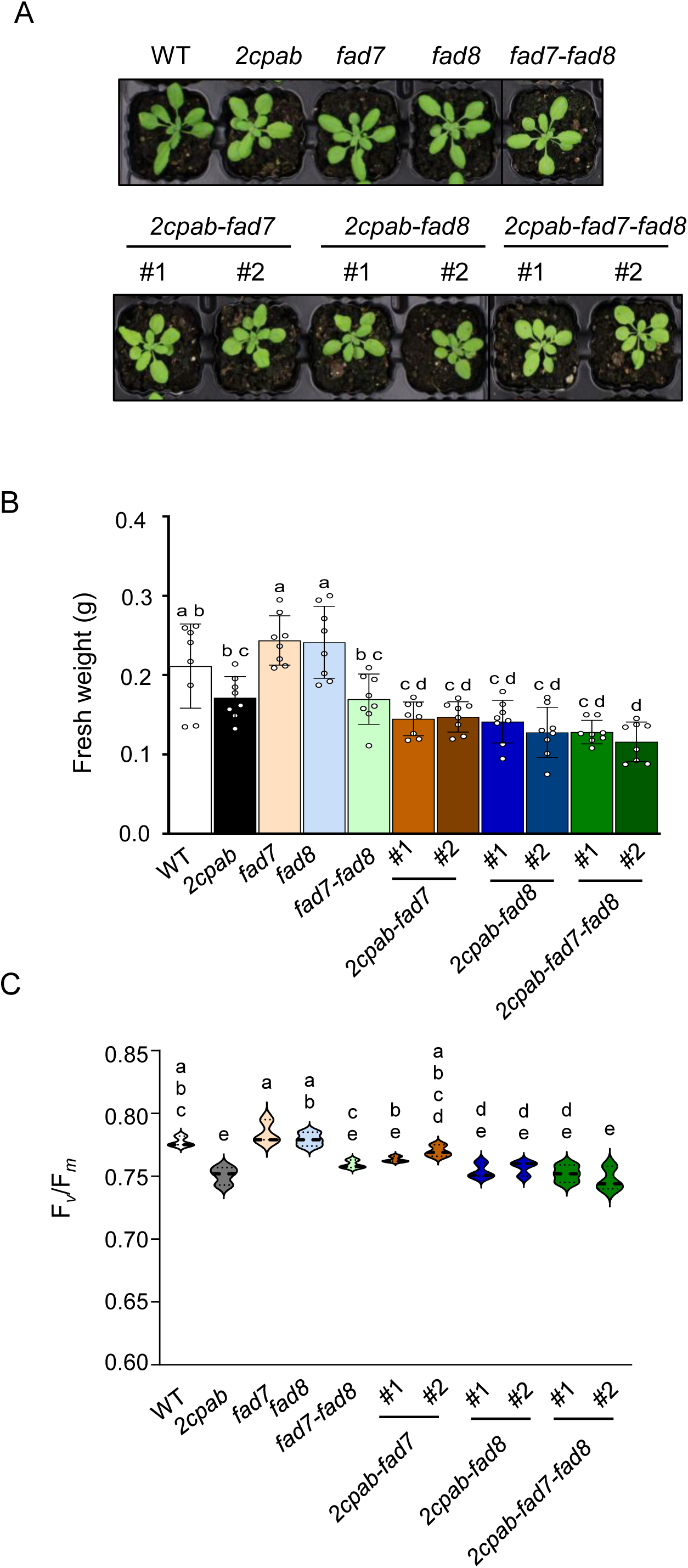
Arabidopsis plants lacking 2-Cys Prxs and chloroplast ω-3 fatty acid desaturases display similar phenotype to that of the *2capb* mutant. **(A)** Plants of the wild-type (WT) and mutant lines, as indicated, grown under long-day condition at a light intensity of 125 μE m^−2^ s^−1^ for 4 weeks. **(B)** Weight of the rosette leaves determined from at least 7 plant grown as (A) and represented as average values ± SD. **(C)** Maximum PSII quantum yield of dark-adapted leaves grown as (A) determined as the ratio of variable fluorescence (F*_v_*) to maximal fluorescence (F*_m_*), F*_v_*/F*_m_*. Data are means ± SD of four independent plants. Letters indicate significantly different according to one-way ANOVA with a Tukey’s multiple comparisons test (*p* < 0.05).

Interestingly, significant differences were observed in the composition of fatty acids in rosette leaves between the multiple *2cpab-fad* mutants and their parental *2cpab* and *fad* mutant lines. Principal component analysis (PCA) of fatty acid proportions (% of total) revealed a clear separation among the different lines along the first two principal components (Fig. 2A). The first principal component (PC 1) explained 97.9% of the total variance, while the second one (PC 2) accounted for 1.6%, together explaining nearly all of the total variance. Only two mutants *2cpab-fad7-fad8* and *fad7-fad8*, clustered together (Fig. 2A), indicating that their fatty acid compositions were similar. The WT, *2cpab*, *fad7*, *fad8*, and *2cpab-fad8* lines were positioned on the positive side of PC 1, whereas *2cpab-fad7* and the cluster formed by *fad7-fad8* and *2cpab-fad7-fad8* appeared on the negative side (Fig. 2A), indicating that these latter groups were the most distinct in their fatty acid composition. This distribution correlated with the levels of TAs (16:3 and 18:3) observed in the different mutant lines (Fig. 2B), which led to several relevant findings. First, the similar levels of 18:3 observed in *2cpab-fad7-fad8,* and *fad7-fad8* mutants explained that these lines clustered together in the PCA. Additionally, these results indicate that the absence of 2-Cys Prxs did not further affect 18:3 production by microsomal ω-3 FAD3, the only functional ω-3 FAD in these mutant lines, nor its subsequent incorporation into chloroplast lipids via the eukaryotic pathway. Second, both chloroplast ω-3 FADs functions were affected by the absence of 2-Cys Prxs since TAs levels in both multiple mutants, *2cpab-fad7* and *2cpab-fad8,* were lower than in their parental *fad7* and *fad8* single mutants, respectively (Fig. 2B). However, the lower TAs levels in the *2cpab-fad7* mutant than in *2cpab-fad8* suggests a more severe impact of the lack of 2-Cys Prxs on FAD8 than on FAD7 activity. Finally, it should be noted that neither of the two chloroplast ω-3 FADs completely losses their activity in the absence of 2-Cys Prxs, as TAs levels were higher in both multiple mutants than in the *fad7-fad8* double mutant (Fig. 2B).

**Figure 2.**
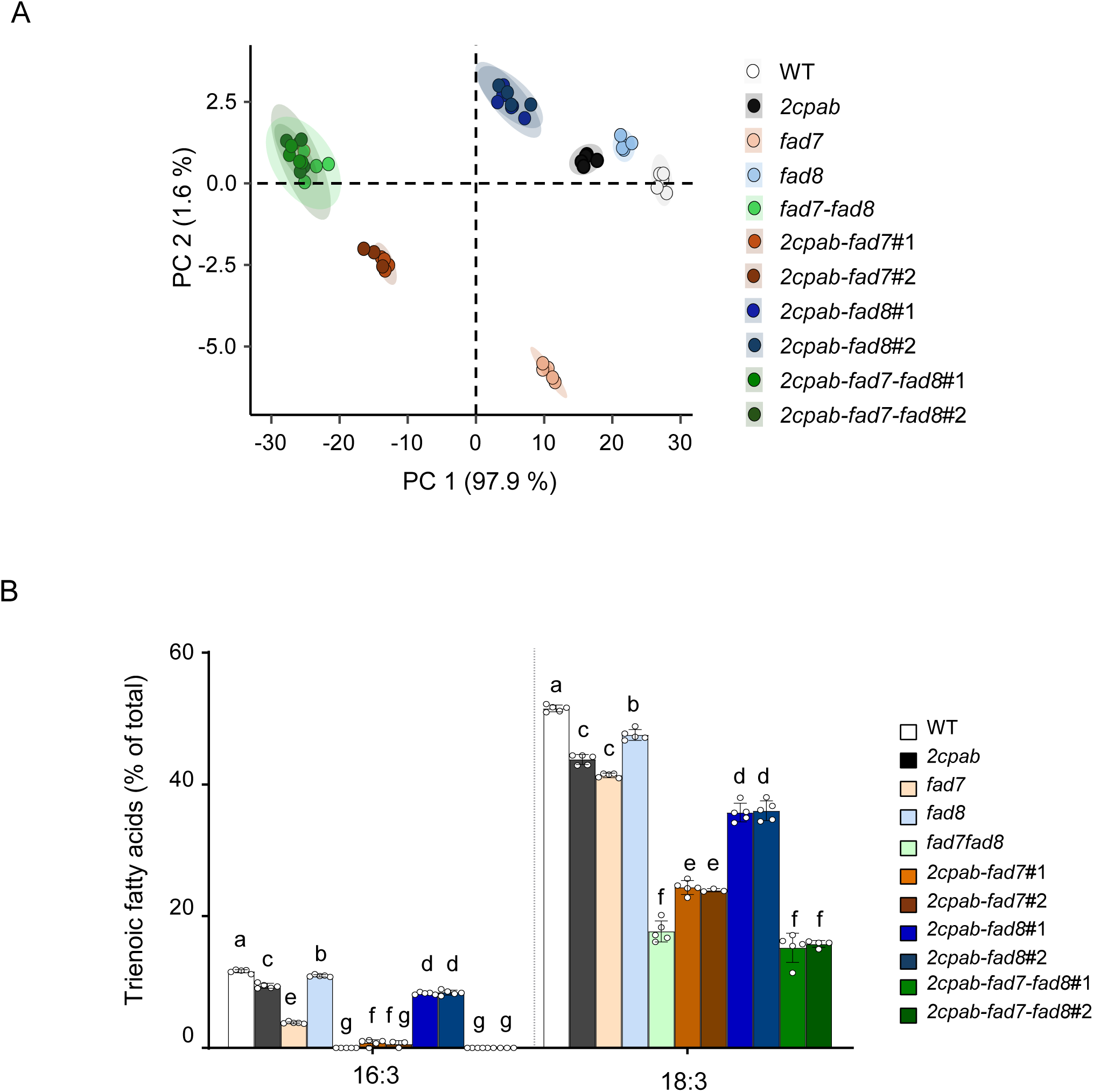
Chloroplast ω-3 fatty acid desaturases FAD7 and FAD8 functions are affected in the absence of 2-Cys Prxs. Wild-type (WT) and mutant lines, as indicated, were grown under long-day conditions at a light intensity of 125 μE m^−2^ s^−1^ for 4 weeks. **(A)** Principal component analysis of leaf total fatty acid composition. Score plot showing the distribution of samples according to the first two principal components (PC 1 and PC 2), which explain 97.9% and 1.6% of the total variance, respectively. Each point represents an independent biological replicate, and colours indicate the different genotypes. **(B)** Trienoic fatty acid levels (16:3 and 18:3) in leaves from the different lines grown as (A). Total leaf lipids were extracted and total fatty acid composition analysed as described in Materials and Methods. Data are means ± SD of five independent plants. Letters indicate significantly different by one-way ANOVA with a Tukey’s multiple comparison test (*p* < 0.05).

To gain further insight into the role of 2-Cys Prxs in the function of chloroplast ω-3 FADs, the fatty acid composition of the different membrane lipid classes was analysed, and the TA ratio between each *2cpab-fad* multiple mutant line versus its corresponding *fad* mutant parental line was determined. Ratios lower than 1 indicate negative effect of the absence of 2-Cys Prxs on the remaining FAD, whereas a ratio of ∼1 indicates no effect. As expected, the 18:3 ratios in the main extra-chloroplast membrane lipids (PI+PS, PE, PA and PC) were close to 1, confirming the minor effect, if any, of 2-Cys Prxs on FAD3 function (Fig. 3A-D). Among the chloroplast lipids, the 18:3 and 16:3 ratios in galactolipids were differently affected, whereas the *2cpab-fad8* lines exhibited 18:3 and 16:3 ratios of ∼0.8, in both galactolipids, a much stronger reduction was observed in *2cpab-fad7* lines, with 18:3 and 16:3 ratios decreasing to ∼0.4 and ∼0.2, respectively, in both MGDG and DGDG (Fig. 3G-J). Thus, these results indicate that 2-Cys Prxs are required for efficient FAD8-dependent desaturation of galactolipid 18:2 and 16:2 fatty acids, whereas FAD7 function is marginally affected. However, the most striking observation was the strong reduction in the 18:3 ratio in PG (∼0.4) in both the *2cpab-fad7* and *2cpab-fad8* mutants (Fig. 3E), indicating a specific role of 2-Cys Prxs in the desaturation of PG-18:2 by the two chloroplast ω-3 FADs.

**Figure 3.**
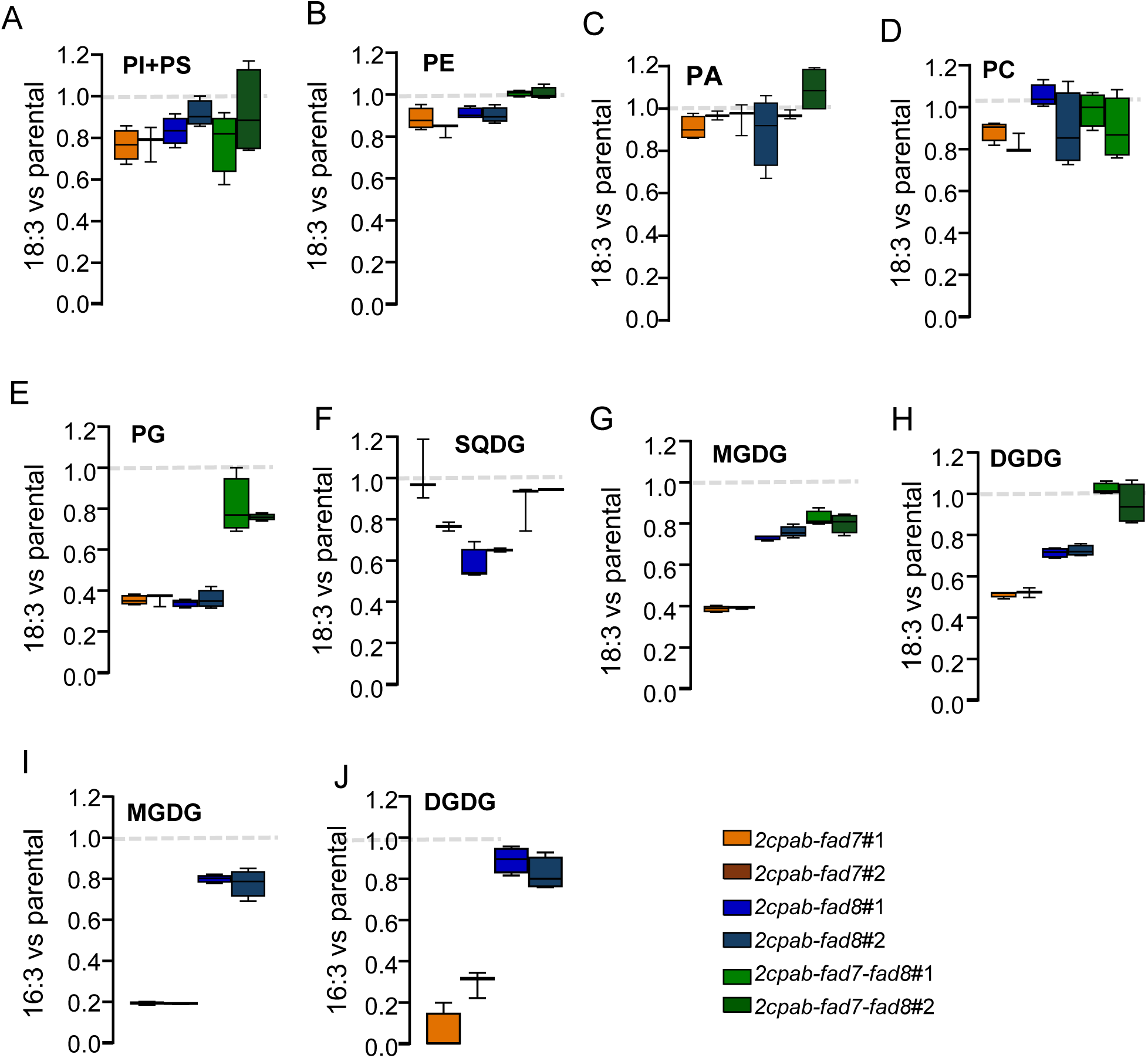
2-Cys Prxs are required for efficient ω-3 desaturation in thylakoid membrane lipids. Wild-type (WT) and mutant lines, as indicated, were grown under long-day conditions at a light intensity of 125 μE m^−2^ s^−1^ for 4 weeks. Total leaf lipids were extracted, the different lipid classes separated by thin layer chromatography (TLC) and fatty acid compositions analysed as described in Materials and Methods. Graphics represent the TA ratio in each *2cpab-fad* multiple mutant line relative to their corresponding *fad* parental mutant. A ratio lower than 1 indicates that the absence of 2-Cys Prxs negatively affects the functional desaturase enzyme in each membrane lipid and multiple mutant line. Data are means ± SD of five independent plants. The 18:3 ratio is represented in **(A)** phosphatidylinositol (PI) + phosphatidylserine (PS), **(B)** phosphatidylethanolamine (PE), **(C)** phosphatidic acid (PA), **(D)** phosphatidylcholine (PC), **(E)** phosphatidylglycerol (PG), **(F)** sulfoquinovosyldiacylglycerol (SQDG), **(G)** monogalactosyldiacylglycerol (MGDG) and **(H)** digalactosyldiacylglcyerol (DGDG). The 16:3 ratio in represented in **(I)** MGDG and **(J)** DGDG.

### 2-Cys Prxs interact with chloroplast ω-3 FAD7 and FAD8

The effect of the lack of 2-Cys Prxs on the two chloroplast ω-3 FADs raises the issue of the molecular basis of the functional relationship between these enzymes. Previous analyses of Arabidopsis lines over-expressing FAD8 in the WT and the *2cpab* genetic backgrounds (WT/FAD8-OE and *2cpab*/FAD8-OE) led to the notion that the effect of 2-Cys Prxs on TAs levels is exerted via the stability of FAD8 (Hernández et al., 2024). Thus, to further analyse the effect of 2-Cys Prxs on fatty acid unsaturation, a similar approach was set out to test the contribution of 2-Cys Prxs to FAD7 activity by over-expressing FAD7 in the WT and the *2cpab* genetic backgrounds. To that end, Arabidopsis WT and *2cpab* mutant plants were transformed with constructs containing the *FAD7* gene, fused to CFP and HA tags at the C-terminus, under the control of the CaMV 35S promoter. Two independent transgenic lines, WT/FAD7-OE and *2cpab*/FAD7-OE, respectively, were selected, and the level of expression of FAD7 was compared with that of FAD8 in the previously reported WT/FAD8-OE and *2cpab*/FAD8-OE (Hernández et al., 2024). Western blot analysis showed FAD7 protein levels comparable to those of FAD8 in both genetic backgrounds, with significantly lower amount of the overexpressed FAD proteins when 2-Cys Prxs are not present (Fig. S3A-B), suggesting similar effects of 2-Cys Prxs on the stability of FAD7 and FAD8.

The contribution of 2-Cys Prxs to the stability of FAD7 and FAD8 might be exerted indirectly, via the effect of the lack of 2-Cys Prxs on the redox state of the chloroplast, or via the direct interaction of these proteins. To test these possibilities, we performed coimmunoprecipitation experiments with the anti-HA antibody. When membrane fractions from formaldehyde-fixed leaves of WT/FAD7-OE plants were used a band corresponding to 2-Cys Prxs was identified in the eluted fraction, which was not present in WT plants (Fig. 4A). Moreover, CHL27, included in these experiments as a negative control, was not immunoprecipitated with the anti-HA antibody in membrane fractions of neither the WT/FAD7-OE nor the WT plants (Fig. 4A). Thus, these experiments show the *in vivo* interaction of 2-Cys Prxs and FAD7, which is in line with the previously reported interaction of 2-Cys Prxs and FAD8 (Hernández et al., 2024). The interaction of 2-Cys Prxs with FAD7 and FAD8 was further tested by an alternative approach, a membrane optimized split-ubiquitin yeast two-hybrid method. To that end, FAD7 and FAD8, lacking their respective transit peptides to avoid potential targeting to yeast mitochondria, were fused with the C-terminal half of ubiquitin and the artificial transcription factor PLV as bait construct, and 2-Cys Prxs A and B were fused with a mutated form of the N-terminal half of ubiquitin, as prey constructs. Both 2-Cys Prxs interacted with FAD7 and FAD8, as demonstrated by yeast growth on histidine-deficient medium, whereas no growth was observed with the bait or prey plasmid alone or when KAT1 was used as a prey control (Fig. 4B).

**Figure 4.**
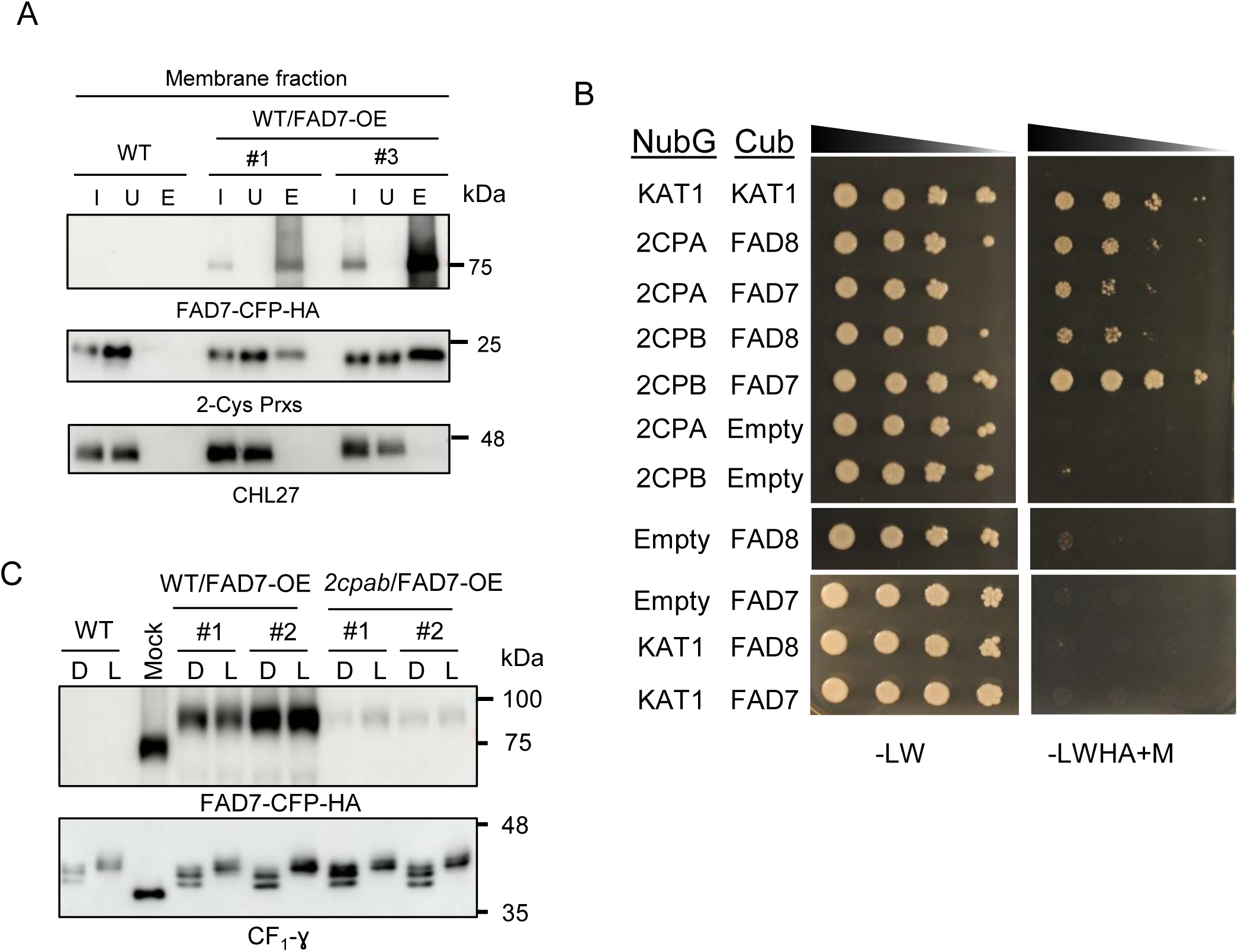
2-Cys Prxs interacts with chloroplast ω-3 fatty acid desaturases FAD7 and FAD8. **(A)** Coimmunoprecipitation assay using membrane fractions obtained from formaldehyde-fixed leaves from WT and WT/FAD7-OE plants, grown under long-day conditions at a light intensity of 125 μE m^−2^ s^−1^ for 4 weeks. Proteins preparations (2 µg) from the input (I) and unbounded (U) fractions and ∼25% of the total eluted volume (E) were immunoblotted and probed with anti-HA, indicating the presence of FAD7-CFP-HA, anti-2-Cys Prxs, and anti-CHL27. Molecular mass markers (kDa) are indicated on the right. **(B)** Split-ubiquitin assay showing interaction of 2-Cys Prxs A and B with FAD7 and FAD8. The prototypic growth assay of yeast containing bait (Cub) and prey (NubG) vectors on non-selective media (-LW) are shown on the left, and selective media (-LWHA+Methionine) on the right. Positive control consisted of KAT1 in the NubG and Cub vectors and negative interaction was demonstrated by empty bait or prey vectors, and KAT1 as prey with FAD7 and FAD8 in the bait vector. (C) *In vivo* redox state of YFP–HA tagged FAD7 in rosette leaves from WT/FAD7-CFP-HA and *2cpab*/FAD7-YFP-HA plants, grown as in (A), determined at the end of the dark period (D), and after 30 min of illumination at 150 μE m^−2^ s^−1^ (L) by labeling of the thiol groups with the alkylating agent MM(PEG)_24_. A non-alkylated WT/FAD7-YFP-HA sample (mock) is shown as a control. Molecular mass markers (kDa) are indicated on the right.

To gain further insight into the potential structural basis of the interaction between the 2-Cys Prx decamer and FAD7 or FAD8, *in silico* structure predictions were performed using AlphaFold 3 (Fig. S4A-B). In a subset of the predicted models, FAD7 and FAD8 were positioned within the central cavity of the 2-Cys Prx decameric ring. However, the confidence metrics for the predicted complexes were relatively low (ipTM and pTM values of approximately 0.35), indicating substantial uncertainty regarding the predicted protein–protein arrangement. This low confidence may partly reflect the limitations of modelling membrane-associated proteins in the absence of a lipid bilayer environment. Nevertheless, the association between 2-Cys Prx and FAD7/FAD8 is supported by our experimental data, as previously demonstrated *in vivo* by co-immunoprecipitation (Fig. 4A) and further supported by a split-ubiquitin assay (Fig. 4B). Although the predicted models only show a possible structural hypothesis for the association between these proteins, we further examined the intermolecular interactions in the representative models shown in Fig. S4A-B. The analysis of the FAD7–2-Cys Prx model identified three intermolecular interactions, including hydrogen bonds and electrostatic interactions (Supplementary Table S4). Among these, two were identified as salt bridges: one between Lys1 of a 2-Cys Prx subunit and Asp18 of FAD7, and another between Lys1 of a 2-Cys Prx subunit and Asp35 of FAD7. Similarly, the analysis of the FAD8–2-Cys Prx model identified five intermolecular interactions, including hydrogen bonds and electrostatic interactions (Supplementary Table S5). Among these, one was identified as a salt bridge between Lys1 of a 2-Cys Prx subunit and Glu383 of FAD8.

Previously, it was shown that the absence of 2-Cys Prxs does not affect the redox state of FAD8, suggesting that the function of 2-Cys Prxs on thylakoid fatty acid unsaturation is exerted beyond the classic redox regulatory mechanism (Hernández et al., 2024). Similarly to FAD8, the FAD7 mature protein contains 4 cysteines. Cys-109 (Cys-102 in FAD8) and Cys-165 (Cys-158 in FAD8) are expected to be localized at hydrophilic regions of the proteins facing the stromal side of the chloroplast inner envelope membrane, while Cys-281 and -291 (Cys-274 and -284 in FAD8) have expected location in a transmembrane domain (Fig. S4C-D). To test whether FAD7 is subjected to redox regulation, thiol-specific labelling with the alkylation agent MM(PEG)_24_ was performed on leaf protein extracts obtained from dark- and light-acclimated WT/FAD7-OE and *2cpab*/FAD7-OE transgenic plants. Immunoblot analysis revealed a single band in both transgenic lines with a shift in the molecular weight due to the presence of the alkylating agent, indicating labelling of Cys-residues in its thiolic form (Fig. 4C). The electrophoretic mobility was similar in all transgenic lines regardless of the genetic background and light condition, in contrast to the γ subunit of ATPase (CF_1_-γ), a well-established redox regulated protein, which is rapidly reduced in the light (Yoshida et al., 2014). Overall, these results indicate that 2-Cys Prxs interact with FAD7 but have no effect on its redox state, as previously reported for FAD8 (Hernández et al., 2024).

### Role of 2-Cys Prxs and PG unsaturation levels in Arabidopsis long-term cold response

The levels of *cis*-unsaturated fatty acids in PG have been associated with the protection of PSI and PSII from photoinhibition, particularly under cold stress (Moon et al., 1995; Sui et al., 2007; Sun et al., 2011; Ivanov et al., 2012). Moreover, it is well stablished that TAs are important for photosynthetic performance at low temperatures (Routaboul et al., 2000; Vijayan and Browse, 2002). Thus, the results showing the interaction between 2-Cys Prxs and chloroplast ω-3 FADs and the role of 2-Cys Prxs controlling the ω-3 desaturation specifically in PG prompted us to investigate the contribution of these enzymes to long-term acclimation to cold. For this purpose, the mutants under analysis were grown under short-day (SD) conditions (8 h light/16 h dark) for 1 month at 22 °C followed by an additional month at 10 °C during the day and 5 °C at night. Cold conditions exerted a more pronounced effect on growth rates in mutant lines lacking 2-Cys Prxs, *2cpab*, *2cpab-fad7*, *2cpab-fad8* and *2cpab-fad7-fad8* than standard SD growth conditions, though no significant differences were detected among the mutant genotypes (Fig. 5A-B). Notably, the *fad7-fad8* double mutant showed a marked growth inhibition under cold conditions (Fig. 5B), accompanied by a significant decline in photosynthetic efficiency, which was also observed in the quadruple mutant *2cpab-fad7-fad8* (Fig.5C). On the contrary, the combined loss of 2-Cys Prxs and FAD7 or FAD8 in the *2cpab-fad7* and *2cpab-fad8* mutants, respectively, barely affected F*_v_*/F*_m_* values under cold stress, which were comparable to those of the *2cpab* mutant and slightly lower than those of the WT (Fig. 5C).

**Figure 5.**
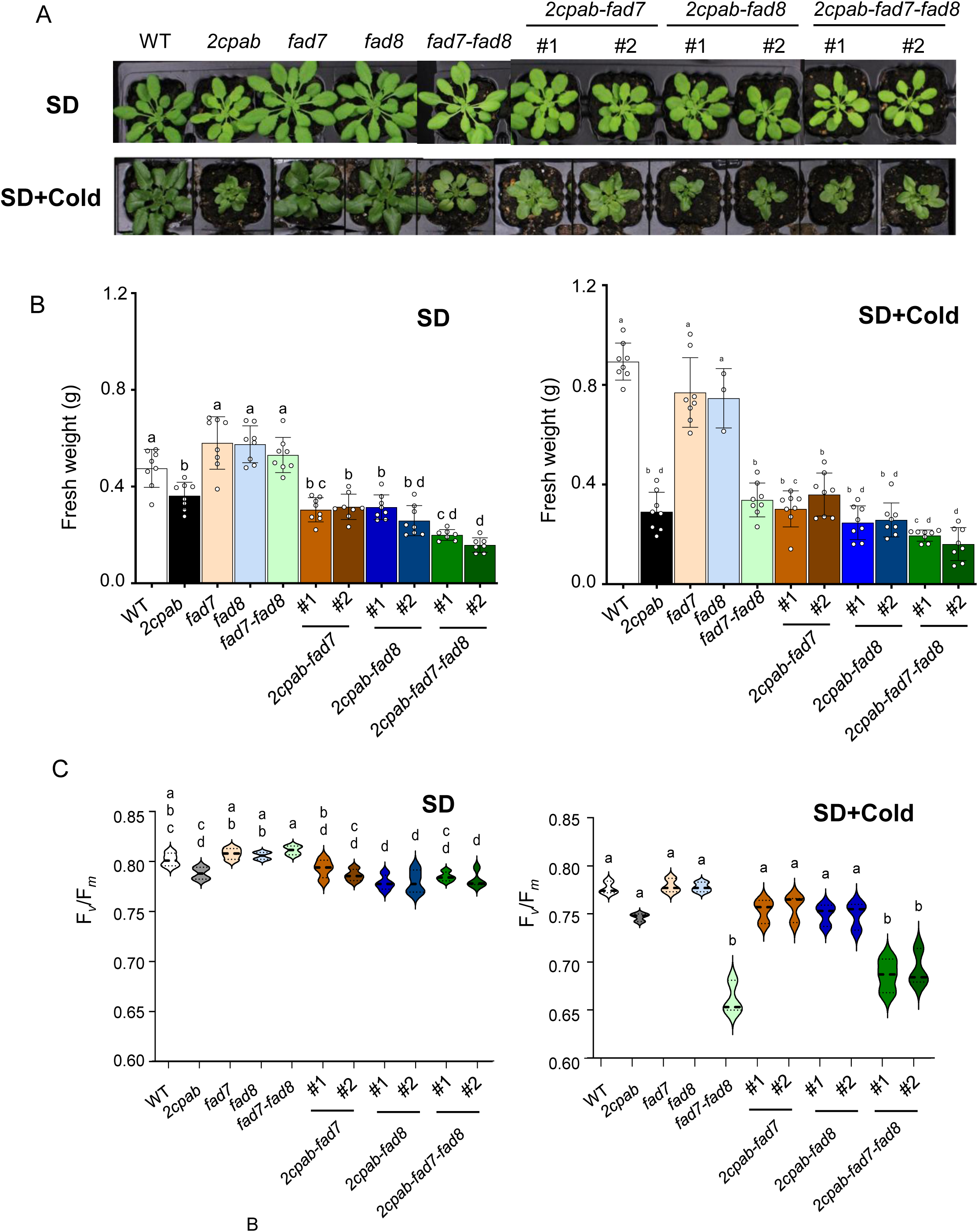
The combined loss of 2-Cys Prxs and chloroplast ω-3 fatty acid desaturases functions barely affect photosynthetic performance under cold conditions. **(A)** Plants of the wild-type (WT) and mutant lines, as indicated, grown under short-day condition at a light intensity of 125 μE m^−2^ s^−1^ (SD) for 45 d, and during 28 d under SD conditions and an additional month with the same photoperiod and light intensity except that temperature was set at 10 °C and 5 °C during light and dark periods, respectively (SD+Cold). **(B)** Weight of the rosette leaves determined from at least 7 plant grown as (A) and represented as average values ± SD. **(C)** Maximum PSII quantum yield of dark-adapted leaves grown as (A) determined as the ratio of variable fluorescence (F*_v_*) to maximal fluorescence (F*_m_*), F*_v_*/F*_m_*. Data are means ± SD of four independent plants. Letters indicate significantly different according to one-way ANOVA with a Tukey’s multiple comparisons test (*p* < 0.05).

PCA of fatty acids proportions (% of total) of all the lines under study revealed that the genetic background, rather than growth conditions, was the primary source of variation (Fig. 6A). However, within each genotype, samples grown under cold conditions (SD+Cold) clustered separately from those grown under control SD conditions, indicating that all lines underwent remodelling of leaf fatty acid composition in response to low temperature. Notably, the WT and *2cpab* mutant clustered together under cold conditions, but not under control conditions (Fig. 6A), as expected since TAs levels in the *2cpab* mutant were like those in WT plants grown under cold conditions (Fig. 6B). It is known that the *FAD8* gene, but not *FAD7*, is induced by cold in Arabidopsis leaves (McConn et al., 1994; Román et al., 2015; Soria-García et al., 2019), consistent with the increase in the unsaturation level of thylakoid membrane lipids to maintain membrane fluidity. Interestingly, mutants lacking 2-Cys Prxs exhibited increased expression of both chloroplasts ω-3 *FAD* genes compared to WT plants (Fig. S5), which could explain the restoration of TAs levels in the *2cpab* mutant to WT values. Consistent with this observation, the triple mutants *2cpab-fad7* and *2cpab-fad8* showed reduced TAs levels compared to the *2cpab* mutant and the WT, as only *FAD8* or *FAD7* transcript levels, respectively, were like those of the *2cpab*. Therefore, it is likely that the induction of both chloroplast ω-3 *FAD* genes is required to maintain TAs levels in thylakoid membranes under cold conditions when 2-Cys Prxs are not present. Additionally, the significantly lower TA levels detected in the *2cpab-fad7* mutant compared to those of *2cpab-fad8* (Fig. 6B), further support the notion that FAD8 function was more severely compromised than FAD7 in the absence of 2-Cys Prxs. Thus, to gain further insight into the contribution of 2-Cys Prxs to TAs composition in response to cold treatment, the ratios of 18:3 and 16:3 TAs in each class of lipids was determined for the different mutants versus their parentals. Although TAs levels increased in the main chloroplast lipids in all genotypes grown in cold in comparison to control grown plants, in the *2cpab-fad7* mutant, the FAD8-dependent desaturation of 18:2- (Fig. 7A-C) and 16:2- (Fi. 7D-E) in the absence of 2-Cys Prxs, was impaired in thylakoid lipids. However, FAD7-dependent desaturation in the *2cpab-fad8* mutant was only affected in PG (Fig. 7). These findings further confirm the role for 2-Cys Prxs in maintaining TA levels specifically in PG. Interestingly, although the absence of 2-Cys Prxs resulted in lower PG-18:3 levels in both triple mutants to approximately 30-50% of WT levels, no significant effect on photosynthetic performance under cold conditions was observed as shown by the comparable F*_v_*/F*_m_* values in the *2cpab-fad7*, *2cpab-fad8*, *2cpab* mutants, and remained higher than those observed in *fad7-fad8* and the quadruple mutant *2cpab-fad7-fad8* (Fig. 5C), both of which exhibited lower TA levels in PG (∼5%).

**Figure 6.**
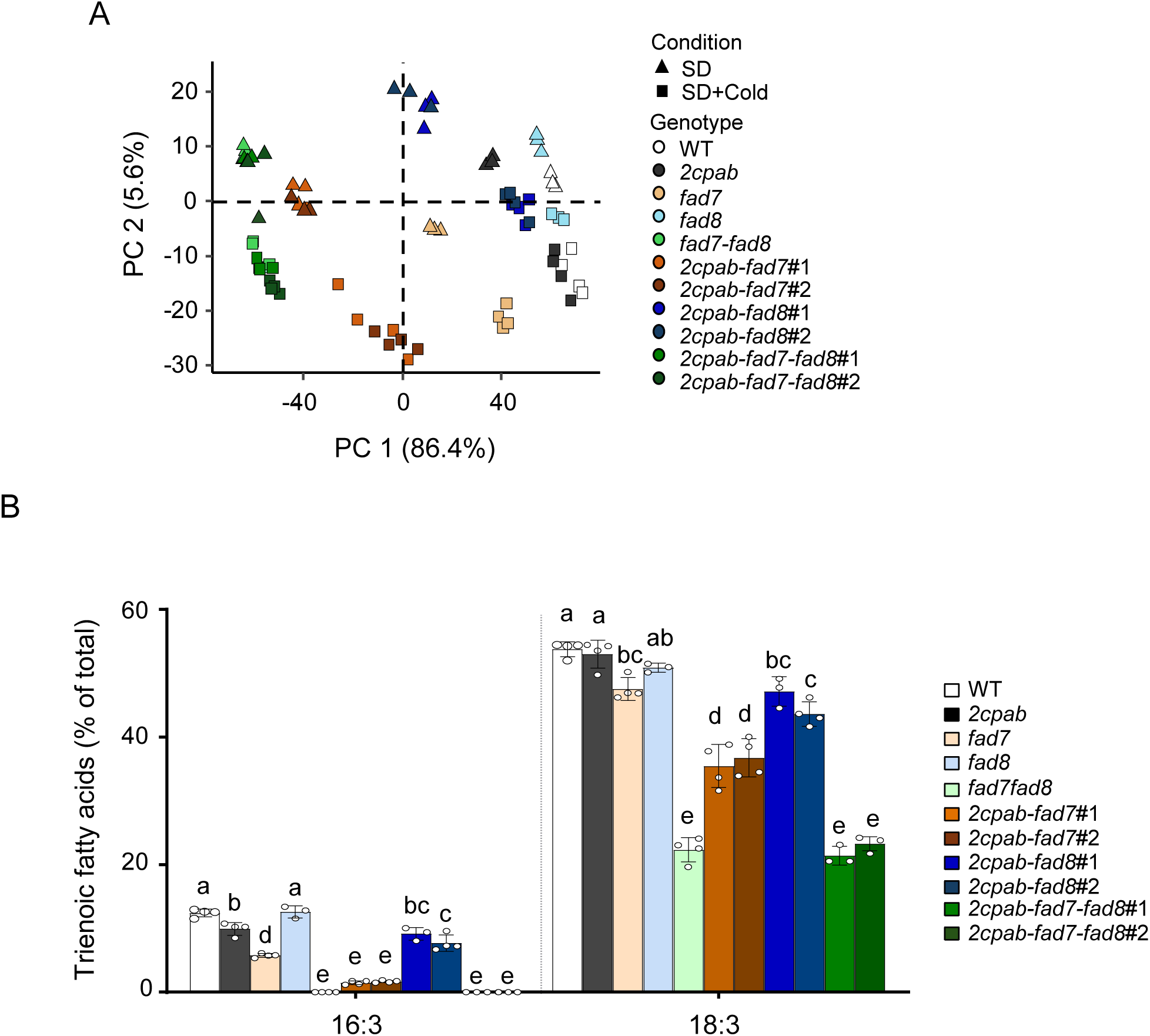
Chloroplast ω-3 fatty acid desaturases FAD7 and FAD8 are required in the absence of 2-Cys Prxs to maintain trienoic fatty acid levels in Arabidopsis leaves under cold conditions. Plants of the wild-type (WT) and mutant lines, as indicated, grown under short-day condition at a light intensity of 125 μE m^−2^ s^−1^ (SD) for 45 d, and during 28 d under SD conditions and an additional month with the same photoperiod and light intensity except that temperature was set at 10 °C and 5 °C during light and dark periods, respectively (SD+Cold). **(A)** Principal component analysis of leaf total fatty acid composition. Score plot showing the distribution of samples according to the first two principal components (PC 1 and PC 2), which explain 86.4% and 5.6% of the total variance, respectively. Each point represents an independent biological replicate. Growth conditions are indicated by symbol shape, whereas genotypes are distinguished by colours. **(B)** Trienoic fatty acid levels (16:3 and 18:3) in leaves from the different genotypes grown under SD+Cold conditions. Total leaf lipids were extracted and total fatty acid composition analysed as described in Materials and Methods. Data are means ± SD of four independent plants. Letters indicate significantly different by one-way ANOVA with a Tukey’s multiple comparison test (*p* < 0.05).

**Figure 7.**
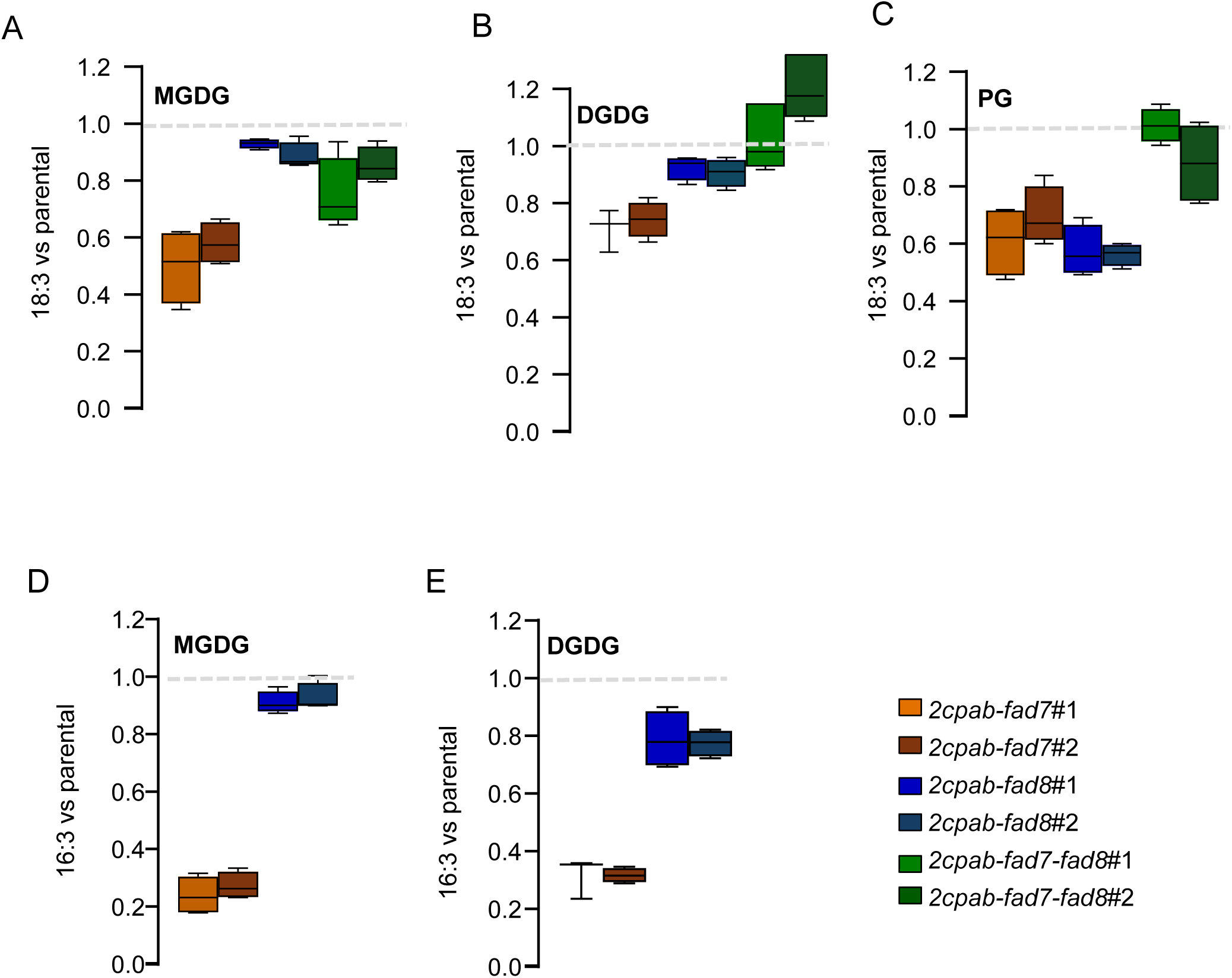
2-Cys Prxs are required for efficient ω-3 desaturation in thylakoid membrane lipids under cold conditions. Plants of the wild-type (WT) and mutant lines, as indicated, grown under short-day condition at a light intensity of 125 μE m^−2^ s^−1^ (SD) for 45 d, and during 28 d under SD conditions and an additional month with the same photoperiod and light intensity except that temperature was set at 10 °C and 5 °C during light and dark periods, respectively (SD+Cold). Total leaf lipids were extracted, the different lipid classes separated by thin layer chromatography (TLC) and fatty acid compositions analysed as described in Materials and Methods. Graphics represent the TA ratio in each *2cpab-fad* multiple mutant line relative to their corresponding *fad* parental mutant. A ratio lower than 1 indicates that the absence of 2-Cys Prxs negatively affects the functional desaturase enzyme in each membrane lipid and multiple mutant line. Data are means ± SD of five independent plants. The 18:3 ratio is represented in **(A)** monogalactosyldiacylglycerol (MGDG), **(B)** digalactosyldiacylglcyerol (DGDG) and **(C)** phosphatidylglycerol (PG). The 16:3 ratio in represented in **(D)** MGDG and **(E)** DGDG.

### The role of 2-Cys Prxs and chloroplast ω-3 FADs in the response to high light

A previous study reported the enhanced sensitivity of the *2cpab* mutant to high light (HL) (Awad et al., 2015). Since the thylakoid membrane fluidity plays a key role in the PSII repair cycle at HL (Yamamoto, 2016; Verhoeven et al., 2026), we analysed the effects of the combined deficiencies of 2-Cys Prxs and chloroplast ω-3 FADs in plant response to this environmental stress. For this purpose, the lines under study, grown under standard conditions at 125 μE m^−2^ s^−1^, were exposed to a light intensity of 900 μE m^−2^ s^−1^ (HL) at 15°C, conditions known to delay the recovery of photodamaged PSII in Arabidopsis (Takahashi and Murata, 2008). During the first 6 h of treatment, all the genotypes exhibited PSII photoinactivation, as evidenced by a rapid reduction in F*_v_*/F*_m_* values, though the effect was more severe in the quadruple mutant *2cpab-fad7-fad8* lines (Fig. S6). After 24 h under HL, most of the *2cpab-fad7-fad8* plants failed to recover and exhibited leaf photobleaching, as indicated by F*_v_*/F*_m_* values approaching zero, while the remaining genotypes maintained the photosynthetic activity (Fig. 8A). Remarkably, TAs levels were substantially lower in the quadruple mutant *2cpab-fad7-fad8* than in the *2cpab* and the triple mutants *2cpab-fad7* and *2cpab-fad8* (Fig. 8B), in line with the higher sensitivity of this mutant to HL. However, it should be noted that though the *fad7-fad8* and *2cpab-fad7-fad8* mutants show different photosynthetic performance (Fig. 8A), their TAs levels are similar (Fig. 8B), suggesting that 2-Cys Prxs contribute to PSII protection through mechanisms additional to their role in sustaining TA synthesis. Collectively, these findings indicate that both, 2-Cys Prxs and chloroplast ω-3 FADs are required for efficient PSII recovery after photoinhibition by HL.

**Figure 8.**
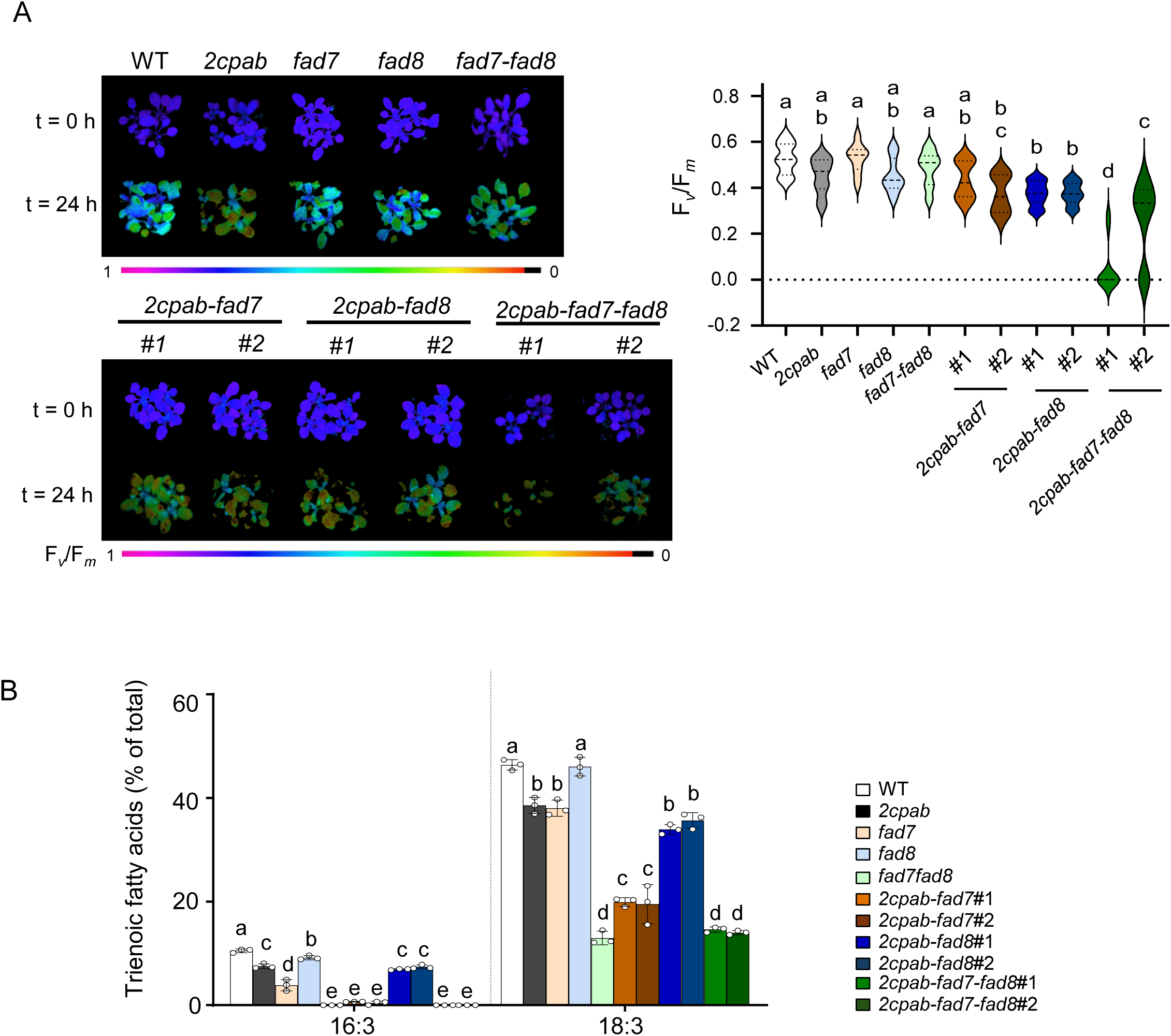
2-Cys Prxs and chloroplast ω-3 fatty acid desaturases cooperate to promote PSII recovery after high-light photoinhibition. Plants of the wild-type (WT) and mutant lines, as indicated, grown under long-day condition at a light intensity of 125 μE m^−2^ s^−1^ for 3 weeks were exposed to a light intensity of 900 µE m^−2^ s^−1^ at 15 °C for 24 h. **(A)** Maximum PSII quantum yield of dark-adapted leaves was represented on the left as colour-images or representative plants of each genotype before and after 24 h of high light treatment and determined on the right as the ratio of variable fluorescence (F*_v_*) to maximal fluorescence (F*_m_*), F*_v_*/F*_m_*. Data are means ± SD of sixteen independent plants. **(B)** Trienoic fatty acid levels (16:3 and 18:3) in leaves from the different genotypes after 24 h of high light treatment. Total leaf lipids were extracted and total fatty acid composition analysed as described in Materials and Methods. Data are means ± SD of three independent plants. Letters indicate significantly different by one-way ANOVA with a Tukey’s multiple comparison test (*p* < 0.05).

## Discussion

### 2-Cys Prxs are required for linolenic acid synthesis in PG

The regulation of fatty acid desaturation, particularly in chloroplast membranes, is a critical mechanism in the response of plants to environmental cues, a process in which the ω-3 desaturases FAD7 and FAD8 of the organelle play key roles. Although in Arabidopsis the *FAD7* gene is constitutively expressed whereas the expression of the *FAD8* gene has been considered stress responsive, a tight coordination of both genes at the transcriptional level is crucial for their specific and non-redundant roles in response to biotic and abiotic stress (Soria-García et al., 2019). In addition, the post-transcriptional control of FAD8 protein stability in response to high temperature has been reported (Matsuda et al., 2005), indicating that FAD8 regulation, at either transcriptional or post-transcriptional levels, could play a significant role in plant response to temperature. However, it is yet unknown the mechanism linking the function of chloroplast ω-3 desaturases to modify the level of TAs in the thylakoid membranes of the organelle in response to environmental stresses. Recently, we found that the stability of FAD8 is compromised in an Arabidopsis mutant devoid of chloroplast 2-Cys Prxs leading to decreased TAs levels in thylakoids lipids (Hernández et al., 2024). These results suggest that 2-Cys Prxs, which are essential in the control of the redox state of the organelle (Pérez-Ruiz et al., 2017; Pérez-Ruiz and Cejudo, 2026), might exert a relevant role in the control of TAs in chloroplast lipids in response to environmental stresses, via the concerted action on FAD7 and FAD8. In this study, we have addressed this issue by using genetic and biochemical approaches to investigate the functional relationship between 2-Cys Prxs and the chloroplast ω-3 FADs. Confirming the previously reported functional relationship between 2-Cys Prxs and FAD8 (Hernández et al., 2024), the desaturation of DA to TA was severely impaired in *2cpab-fad7* mutant lines (Fig. 2B) in the major thylakoid lipids (Fig. 3E-J). Interestingly, whereas FAD7-dependent desaturation in galactolipids was only slightly affected by the loss of 2-Cys Prxs in the *2cpab-fad8* mutant (Fig. 3G-J), the conversion of 18:2 to 18:3 in PG was strongly compromised (Fig. 3E). Overall, these findings reveal the contribution of 2-Cys Prxs to the function of the two chloroplast ω-3 FADs and, most importantly, uncover a relevant role of 2-Cys Prxs in controlling 18:3 levels specifically in PG.

### 2-Cys Prxs may act as scaffold proteins for chloroplast ω-3 desaturation in PG

Several proteins involved in redox homeostasis have previously been implicated in the regulation of fatty acid desaturation in plants. Outside the chloroplast, Prx IIE negatively regulates *FAD3* gene expression during embryo development by promoting the denitrosylation of the transcription factor bZIP67, which decreases its activity thereby reducing *FAD3* transcription (Sánchez-Vicente et al., 2024). Within the chloroplast, the formation of the unique chloroplast fatty acid 16:1^Δ3^ *^trans^* at the *sn*-2 position of PG has also been linked to the activity of redox-related proteins. In this regard, it was shown that FAD4-dependent synthesis of 16:1^Δ3^ *^trans^* requires the presence of Prx Q in Arabidopsis leaves (Horn et al., 2020). Moreover, co-expression of FAD4 and Low Carbon Inducible 2 (LCI2) recruits stromal ascorbate peroxidase (APX) to the vicinity of FAD4 in *Chlamydomonas* thylakoid membranes (Nicodemus et al., 2025). However, the molecular basis of this regulation remains unclear. Prx Q does not appear to alter the redox state of FAD4 (Horn et al., 2020), suggesting that its regulatory effect may be indirect. In line with these observations, the redox state of cysteines in FAD8 (Hernández et al., 2024) and FAD7 (Fig. 4C) were not altered by the content of 2-Cys Prxs. These results, together with the fact that the function of 2-Cys Prxs regulating thylakoid lipids unsaturation level is exerted independently of NTRC (Hernández et al., 2024), the main reductant of theses enzymes (Pérez-Ruiz et al., 2017), indicates that the role of 2-Cys Prxs on TAs synthesis is not exerted by cysteine-based redox exchange reactions.

The fact that 2-Cys Prxs interact with chloroplast ω-3 FADs (Hernández et al., 2024) (Fig. 4A-B), suggests that the molecular basis of this regulation may be related to the ATP-independent holdase or chaperone activity, which is displayed upon the oligomerization into high-molecular weight (HMW) complexes of the enzyme (Ulrich et al., 2021). The mechanism by which 2-Cys Prxs switch from peroxidase to chaperone activity is not yet fully understood. *In vitro* studies have shown that the formation of HMW complexes of 2-Cys Prxs is triggered by the overoxidation of the peroxidatic Cys (Cys_P_) at the active site of these enzymes (Jang et al., 2004; Kumsta and Jakob, 2009; König et al., 2013). In plant 2-Cys Prxs, Mg-ATP or Mg-ADP also induce the formation of HMW complexes that exhibit little or no peroxidase activity but retain moderate chaperone activity (Aran et al., 2008; Aran et al., 2011). In addition, mammalian 2-Cys Prxs form the chaperone-active HMW complexes independently to the overoxidations of the Cys_P_, in response to heat-shock or site-specific phosphorylation (Jang et al., 2006; Saccoccia et al., 2012; Teixeira et al., 2015). Interestingly, the binding of negatively charged lipids, like PS and PG are essential for the formation of human 2-Cys PrxII HMW complexes that display null peroxidase and full chaperone activity (Haruyama et al., 2018). Therefore, considering our results showing a specific role of 2-Cys Prxs in the control of PG TAs, it is tempting to speculate that the association with PG in thylakoid membranes promotes the formation of spherical HMW 2-Cys Prxs complexes that interact with chloroplast ω-3 FADs (Fig. S4A-B), located in the chloroplast envelope membrane (Froehlich et al., 2003; Román et al., 2015). Thus, the 2-Cys Prxs may function as scaffold proteins promoting the spatial association between chloroplast ω-3 FADs and their substrate PG in thylakoid membranes. Consistent with this hypothesis, 2-Cys Prxs have been found to be enriched in the intermediate-density population of proteins proposed to constitute thylakoid-inner envelope contact sites (TIES) in pea (LaBrant et al., 2018). Overall, these results suggest that these proteins may be components of the membrane contact sites, in agreement with previous studies in mice tissues, where 2-Cys Prx was identified in mitochondria-associated endoplasmic reticulum membranes (Wang et al., 2018). Proteins localised in TIES are suggested to represent a functionally specialized domain, acting as ATP-independent chaperones, scaffold proteins or lipid transfer proteins (LaBrant et al., 2026). Remarkably, the TIES proteins identified in this study showed strong co-localization with chlorophyll autofluorescence and exhibited a non-uniform distribution within chloroplast, forming clear punctate structures. Similar localization patterns have been observed for 2-Cys Prxs (Gallardo-Martinez et al., 2023; Rodriguez-Marin et al., 2026) and FAD8 (Román et al., 2015; Hernández et al., 2024). These observations agree with recent studies in *Chlamydomonas* describing the presence of functionally specialized regions within the chloroplast (Wang et al., 2023; Sun et al., 2025) and suggest a role for 2-Cys Prxs in the remodelling of chloroplast membrane unsaturation.

### Concerted roles of 2-Cys Prxs and chloroplast ω-3 FADs in response to long- and short-term PSII photoinhibition

Thylakoid membrane fluidity is a key determinant of plant acclimation to environmental stress, contributing to the tolerance of PSII to photodamage. Under low-temperature conditions, light-induced PSII photoinhibition is exacerbated when the unsaturation of thylakoid membrane lipids is reduced, leading to decreased membrane fluidity, impaired D1 repair, and inefficient PSII reassembly (Moon et al., 1995; Vijayan and Browse, 2002). We have addressed the role of 2-Cys Prxs in the regulation of thylakoid membrane unsaturation degree by challenging mutant plants combining the absence of 2-Cys Prxs and FAD7 and/or FAD8 to prolonged cold (10 °C during the day and 5 °C at night). Interestingly, TAs levels in the *2cpab* mutant were like those of the WT when grown under cold conditions (Fig. 6B). This result correlated with the induction of *FAD7* and *FAD8* genes expression in the *2cpab* mutant compared to the WT (Fig. S5). Although only *FAD8* has been described as cold inducible gene (McConn et al., 1994; Román et al., 2015; Soria-García et al., 2019), increased TAs production by FAD7 has also been reported to contribute to plant acclimation to low, non-freezing temperatures (Kodama et al., 1995; Domínguez et al., 2010). This notion was further supported by the fact that TAs levels in *2cpab-fad7* and *2cpab-fad8* triple mutants were lower than those in the *2cpab* mutant (Fig. 6B), indicating that both chloroplast ω-3 FADs functions are required to maintain WT levels of TAs in plants grown under cold conditions when 2-Cys Prxs are not present. As observed under standard growth conditions, FAD8-dependent desaturation was more severely impaired in any of the chloroplast lipids in the absence of 2-Cys Prxs, whereas the effect on FAD7-dependent desaturation was largely restricted to PG (Fig. 3 and Fig. 7), further confirming the relationship between 2-Cys Prxs and PG unsaturation levels. However, this reduction in TAs levels, particularly in PG, did not affect the photosynthetic performance of plants combining the lack of 2-Cys Prxs and one the chloroplast ω-3 FADs under cold conditions, as indicated by the similar F*_v_*/F*_m_* values observed in *2cpab* and the triple mutants (Fig. 5C). Remarkably, the *fad7-fad8* and the quadruple *2cpab-fad7-fad8* mutants, in which TA levels were even lower (Fig. 6B) and showed minor F*_v_*/F*_m_* values (Fig. 5C), were able to grow under cold conditions (Fig. 5A). These results are consistent with those reported by (Routaboul et al., 2000) showing that 15-20% of the TAs levels of WT plants are sufficient to maintain photosynthetic performance during prolonged exposure to 4 °C. Furthermore, the similar photosynthetic performance of *fad7-fad8* and *2cpab-fad7-fad8* under cold conditions (Fig. 5C) suggests that 2-Cys Prxs do not play a major role in cold acclimation under the conditions tested.

HL is an environmental stress that requires a rapid acclimation response. Excessive irradiance can overwhelm the photosynthetic machinery in chloroplasts, leading to photoinhibition and the generation of reactive oxygen species (ROS). To prevent damage, chloroplasts rapidly activate protective mechanisms such as non-photochemical quenching (NPQ), antioxidant defences, chloroplast movement, and repair of PSII (Khan et al., 2025). When light intensity increases further, the efficiency of photosynthesis, as measured by F*_v_*/F*_m_*, declines significantly. At this stage, the extent of photoinhibition is determined by the balance between light-induced damage and the repair of PSII through D1 protein turnover (Aro et al., 1993), which depends on the fluidity of thylakoid membranes (Yamamoto, 2016; Verhoeven et al., 2026). Lower F*_v_*/F*_m_* recovery in the quadruple mutant *2cpab-fad7-fad8* were observed when plants were exposed for 24 h to HL at 15 °C, compared to the triple mutants *2cpab-fad7* and *2cpab-fad8* and their parental lines (Fig. 8A). Interestingly, TAs levels in this mutant were like those of the *fad7-fad8* (Fig. 8B), meaning that not only membrane fluidity but also additional roles of 2-Cys-Prxs are important for PSII recovery and avoidance of photoinhibition. One possible explanation could be the lower antioxidant capacity of plants lacking 2-Cys Prxs, which has previously been described as HL-sensitive (Awad et al., 2015). HL induces irreversible aggregation of PSII core proteins through ROS and lipid peroxidation, and these aggregates hinder PSII mobility and D1 repair (Yamamoto et al., 2013). Therefore, the combined loss of antioxidant function of 2-Cys Prxs, which detoxify H_2_O_2_ (König et al., 2002) and linoleic acid hydroperoxide (König et al., 2003), together with the decrease in membrane fluidity resulting from the loss of chloroplast ω-3 FADs activity in the *2cpab-fad7-fad8* mutant, may lead to irreversible photoinhibition under HL conditions. An alternative possibility is that the ATP-independent chaperone or holdase activity of 2-Cys Prxs contributes to prevent the irreversible aggregation of PSII core proteins, a process aggravated by reduced thylakoid membrane fluidity, which leads to PSII dysfunction (Yamamoto et al., 2014). The recent funding that 2-Cys Prxs are enriched in discrete chloroplast subdomains that also contain the FAD7 and FAD8 desaturases and the metalloprotease FtsH2 (LaBrant et al. 2026) supports the notion that these enzymes likely participate in processes related to membrane remodelling, that are required for photosynthetic performance and acclimation to stress response. FtsH plays a pivotal role in protein quality control at thylakoid membranes and, accordingly, the lack of FtsH2 in the *var2* mutant results in the accumulation of damaged chloroplast proteins and triggers and UPR-like response, i.e. the accumulation of proteins involved in chloroplast protein quality control (Dogra et al., 2019). The activity of FtsH2 is crucial for extracting substrates from highly viscous thylakoid membranes under cold stress (Zhang et al., 2025), highlighting the close interplay between chloroplast proteostasis and membrane fluidity during the repair of photosynthetic apparatus. Notably, FtsH oligomerization and its proteolytic activity is under redox control in *Chlamydomonas reinhardtii* (Wang et al., 2017) and 2-Cys Prxs are upregulated in the *var2* mutant (Dogra et al., 2019). Overall, these findings open a new research avenue on the role of 2-Cys Prxs in membrane dynamics in response to stress that deserve further investigations.

## Acknowledgment

We are grateful to Carlos Parejo and Alba María Jiménez Segovia (Chromatography Facility) at Instituto de Bioquímica Vegetal y Fotosíntesis (IBVF, Seville, Spain) for technical assistance. We thank Dr. Miguel Alfonso Lozaono (EEAD, Zaragoza, Spain) and Dr. Javier Quintero Toscano (IBVF, Seville, Spain) for providing fatty acid desaturase mutants and split-ubiquitin system vectors and yeast strain, respectively.

## Author contributions

M.L.H., F.J.C., and J.M.P.-R. conceived and designed the study. M.L.H., A.G-R, J.J-L and J.M.P-R generate and characterize mutant and transgenic lines. M.L.H., J.J-L and J.M.P-R performed cold and HL experiments. J.J.-L. conducted the Co-IP and alkylation assays. F.R-M. performed the split-ubiquitin assay. V.O. performed AlphaFold-based prediction of protein interactions. J.M.P-R. conducted RNA extraction and RT-qPCR. M.L.H., J.M.P.-R and F.R-M. analysed the data. M.L.H. wrote the manuscript, with contributions from all authors.

## Supplementary Material

**Supplemental Table S1**. *Arabidopsis thaliana* mutant lines used in this study.

**Supplemental Table S2.** *Arabidopsis thaliana* transgenic lines used in this study.

**Supplemental Table S3.** *Arabidopsis thaliana* oligonucleotides used in this study.

**Supplemental Table S4**. Inter-subunit hydrogen bonds and salt bridges calculated between FAD7 and 2-Cys Prx decamer.

**Supplemental Table S5**. Inter-subunit hydrogen bonds and salt bridges calculated between FAD8 and 2-Cys Prx decamer.

**Supplemental Figure S1.** CRISPR-based editing events of the 2-Cys Prxs genes in Arabidopsis mutant lines lacking 2-Cys Prxs and chloroplast ω-3 fatty acid desaturases.

**Supplemental Figure S2.** Genotypic characterization of chloroplast ω-3 fatty acid desaturases genes in Arabidopsis mutant lines lacking 2-Cys Prxs and chloroplast ω-3 fatty acid desaturases.

**Supplemental Figure S3.** 2-Cys Prxs protein levels in Arabidopsis leaves of transgenic lines overexpressing FAD7 and FAD8 in the WT and 2cpab backgrounds.

**Supplemental Figure S4.** Structural modelling of chloroplast ω-3 fatty acid desaturases FAD7 and FAD8 and its interaction with 2-Cys Prxs using AlphaFold.

**Supplemental Figure S5.** RT-qPCR expression analysis of chloroplast ω-3 desaturases in Arabidopsis wild type and mutant lines lacking 2-Cys Prxs and chloroplast ω-3 fatty acid desaturases.

**Supplemental Figure S6.** Effect of high light on photosynthetic performance in Arabidopsis leaves from WT and mutants lacking 2-Cys Prxs and/or chloroplast ω-3 fatty acid desaturases.

## Data availability statement

All data that support the findings in this paper are available within the article and its Supporting Information or are available from the corresponding authors upon reasonable request.

