## Supplemental Data for "2-Cys peroxiredoxins facilitate phosphatidylglycerol unsaturation and contribute to PSII recovery from high-light photoinhibition"

**Supplemental Table S1**. *Arabidopsis thaliana* mutant lines used in this study.

| Mutant line | Mutation | | | Reference |
|---|---|---|---|---|
| | T-DNA | CRISPR/Cas9 | EMS | |
| *2cpa* | 2-Cys Prx A (SALK_017213) | - | - | Kirchsteiger et al. 2009 |
| *2cpab* | 2-Cys Prx A (GABI_295C05) 2-Cys Prx B (SALK_017213) | - | - | Ojeda et al. 2018 |
| *fad7* | FAD7 (SALK_147096C) | - | - | Román et al. 2015 |
| *fad8* | FAD8 (SALK_093590) | - | - | Román et al. 2015 |
| *fad7-fad8* | - | - | FAD7 (G422A) FAD8 (G446A) | McConn et al. 1994 |
| *2cpab-fad7#1, #2* | FAD7 (SALK_147096C) | 2-Cys Prx A (At3g11630) 2-Cys Prx B (At5g06290) | - | This study |
| *2cpab-fad8#1, #2* | 2-Cys Prx A (GK_295C05) FAD8 (SALK_093590) | 2-Cys Prx B (At5g06290) | | This study |
| *2cpab-fad7-fad8#1* | | 2-Cys Prx A (At3g11630) 2-Cys Prx B (At5g06290) | FAD7 (G141A) FAD8 (G149A) | This study |
| *2cpab-fad7-fad8#2* | FAD7 (SALK_147096C) FAD8 (SALK_093590) | 2-Cys Prx A (At3g11630) 2-Cys Prx B (At5g06290) | | This study |

**Supplemental Table S2.** *Arabidopsis thaliana* transgenic lines used in this study.

| Transgenic line | Background | Promotor | Protein | Reference |
|---|---|---|---|---|
| WT/FAD7-OE | Wild-type | 35S | FAD7-CFP-HA | This study |
| *2cpab*/FAD7-OE | *2cpab* | 35S | FAD7-CFP-HA | This study |
| WT/FAD8-OE | Wild-type | 35S | FAD8-CFP-HA | Hernández et al. 2024 |
| *2cpab*/FAD8-OE | *2cpab* | 35S | FAD8-CFP-HA | Hernández et al. 2024 |

**Supplemental Table S3**. *Arabidopsis thaliana* oligonucleotides used in this study.

| **Genotyping T-DNA primers** | **Locus** | **Sequence** |
|---|---|---|
| ifad7-LP7 | At3g11170 | 5’-ATACAACATCTCCTTCGTCGC-3’ |
| ifad7-RP7 | At3g11170 | 5’-GCTGCTCTGCTTGTTTGTCTC-3’ |
| ifad8-LP8 | At5g05580 | 5’-CTTGGGAAGAGGCTCCAAT-3’ |
| ifad8-RP8 | At5g05580 | 5’-GTAACTTGTCTGCCTGGTTCAC-3’ |
| LBb1.3 | T-DNA | 5’-ATTTTGCCGATTTCGGAAC-3’ |
| **Genotyping CRISPR primers** | **Locus** | **Sequence** |
| 2CPA-F | At3g11630 | 5’-CAACTACTCTCATCTCTTCT-3’ |
| 2CPA-R | At3g11630 | 5’- AGAAGCAGCAACTGTGATTGA -3’ |
| 2CPB-F | At5g06290 | 5’-CGTGGGTCCAAACAGACAGA-3’ |
| 2CPB-R | At5g06290 | 5’- CCGAGGTTGTTGATGGTGGA-3’ |
| **Cloning primers** | **Locus** | **Sequence** |
| attB1-FAD7-g-for | At3g11170 | 5’-AAAAAGCAGGCTtcATGGCGAACTTGGTCTTA-3’ |
| attB2-FAD7-g-rev | At3g11170 | 5’-AGAAAGCTGGGTcATCTGCTCTTACTTTGAC-3’ |
| **Split-Ubiquitin primers** | **Locus** | **Sequence** |
| attB1-2CPA-cds-for | At3g11630 | 5’-GCAGGCTctCCAACCACCATGAAAGCCCAGGCCGAT-3’ |
| attB2-2CPA-cds-rev | At3g11630 | 5’- AGAAAGCTGGGTaAATAGCTGAGAAGTACTCTTTGC-3’ |
| attB1-2CPB-cds-for | At5g06290 | 5’-GCAGGCTctCCAACCACCATGAAGGCTCAGGCTGATGA-3’ |
| attB2-2CPB-cds-rev | At5g06290 | 5’- AGAAAGCTGGGTaGATAGCTGAAAAGTATTCTTTGC-3’ |
| attB1-FAD7-cds-for | At3g11170 | 5’-GCAGGCTctCCAACCACCATGGTGAGCACACCATTAACG-3’ |
| attB2-FAD7-cds-rev | At3g11170 | 5’- AGAAAGCTGGGTaATCTGCTCTTACTTTGACCTC-3’ |
| attB1-FAD8-cds-for | At5g05580 | 5’-GCAGGCTctCCAACCACCATGCCTTCTTCTCTTCTCAATTCC-3’ |
| attB2-FAD8-cds-rev | At5g05580 | 5’- AGAAAGCTGGGTaTGTTCTTTGTCCATTGAGTTTTG-3’ |
| **qRT-PCR primers** | **Locus** | **Sequence** |
| *FAD7* | At3g11170 | 5´-ATGGGTTCACGAGGAATTGG-3´<br>5´-GTGTCGTTAATGGTGTGCTC-3´ |
| *FAD8* | At5g05580 | 5´-TTGCCTCTAACCCTAAACCC-3´<br>5´-GGGAATTGAGAAGAGAAGAAGGA-3´ |
| *ACTIN2* | At3g18780 | 5´-GCACTTGCACCAAGCAGCAT-3´<br>5´-CCTTTCAGGTGGTGCAACGAC-3´ |
| *UBQ10* | At4g05320 | 5´-GGCCTTGTATAATCCCTGATGAATAAG-3´<br>5´-AAAGAGATAACAGGAACGGAAACATAGT-3´ |

The attB recombination sites are underlined. Lower case letters indicate additional nucleotides included to maintain the proper reading frame.

**Supplemental Table S4.** Inter-subunit hydrogen bonds and salt bridges calculated between FAD7 and 2-Cys Prx decamer.

| 2-Cys Prx subunit | 2-Cys Prx Residue | FAD7 subunit | FAD7 Residue | Distance (Å) | Interaction type |
|---|---|---|---|---|---|
| C | Lys 1 | K | Asp 18 | 3.32 | Salt bridge |
| J | Lys 1 | K | Ala 34 | 2.68 | Sidechain-to-backbone H-bond |
| J | Lys 1 | K | Asp 35 | 2.92 | Salt bridge |

**Supplemental Table S5.** Inter-subunit hydrogen bonds and salt bridges calculated between FAD8 and 2-Cys Prx decamer.

| 2-Cys Prx subunit | 2-Cys Prx Residue | FAD8 subunit | FAD78 Residue | Distance (Å) | Interaction type |
|---|---|---|---|---|---|
| B | Lys 1 | K | Tyr 381 | 3.48 | Sidechain-to-backbone H-bond |
| B | Lys 1 | K | Glu 383 | 3.50 | Salt bridge |
| G | Lys 1 | K | Gln 29 | 2.57 | Backbone H-bond |
| I | Gln 3 | K | Thr 36 | 2.01 | Sidechain-to-backbone H-bond |
| C | Gln 3 | K | Lys 288 | 2.48 | Sidechain-to-sidechain H-bond |

### Figure S1

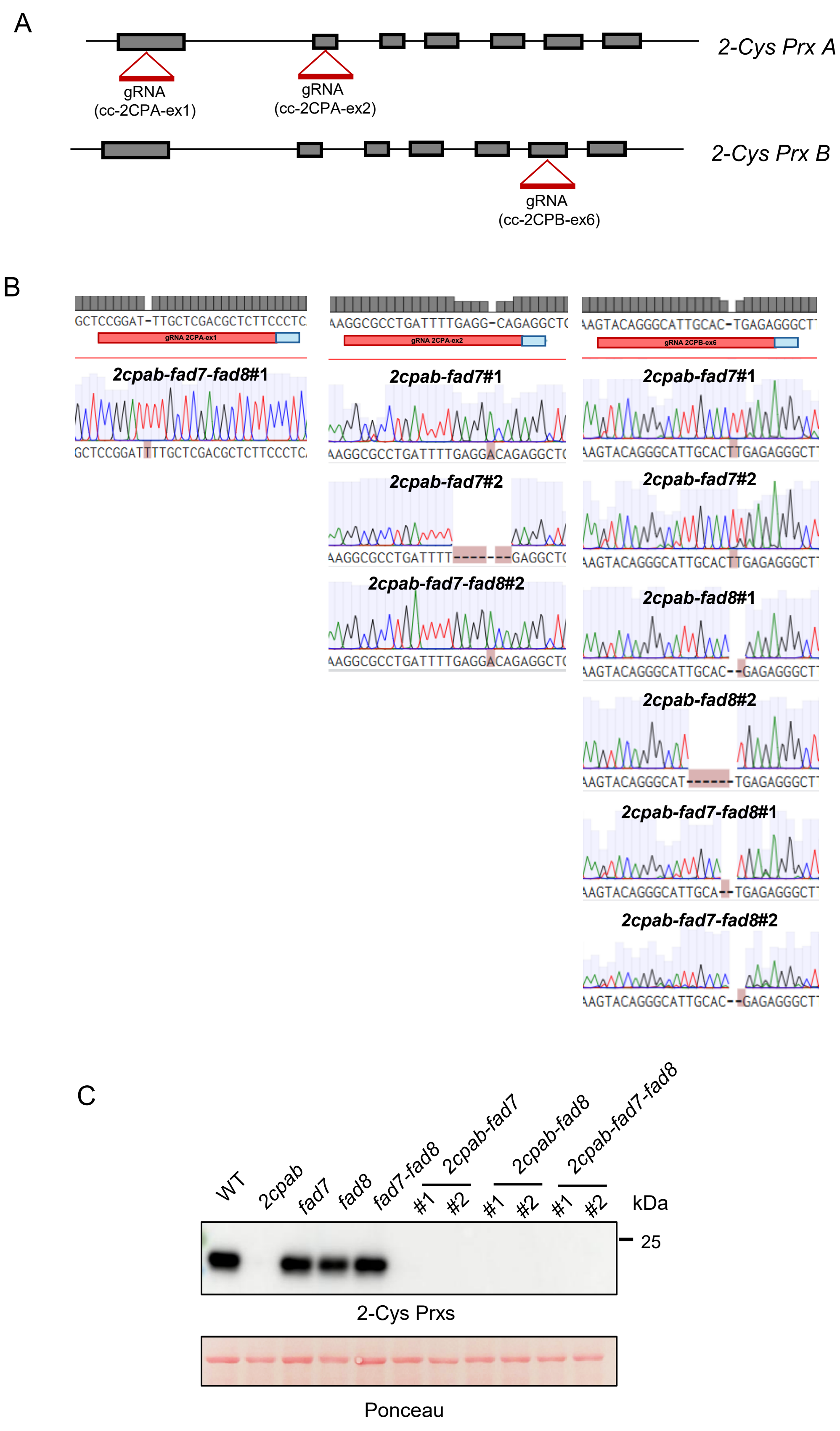

A
2-Cys Prx A
gRNA
(cc-2CPA-ex1)
gRNA
(cc-2CPA-ex2)
2-Cys Prx B
gRNA
(cc-2CPB-ex6)
B
GCTCCGGAT-TTGCTCGACGCTCTTCCCTC
2cpab-fad7-fad8#1
GCTCCGGATTTTGCTCGACGCTCTTCCCTC
AAGGCGCCTGATTTTGAGG-CAGAGGCTC
2cpab-fad7#1
AAGGCGCCTGATTTTGAGGACAGAGGCTC
2cpab-fad7#2
AAGGCGCCTGATTTT------GAGGCTC
2cpab-fad7-fad8#2
AAGGCGCCTGATTTTGAGGACAGAGGCTC
AAGTACAGGGCATTGCAC-TGAGAGGGCT
2cpab-fad7#1
AAGTACAGGGCATTGCACTTGAGAGGGCT
2cpab-fad7#2
AAGTACAGGGCATTGCACTTGAGAGGGCT
2cpab-fad8#1
AAGTACAGGGCATTGCAC--GAGAGGGCT
2cpab-fad8#2
AAGTACAGGGCAT------TGAGAGGGCT
2cpab-fad7-fad8#1
AAGTACAGGGCATTGCA--TGAGAGGGCT
2cpab-fad7-fad8#2
AAGTACAGGGCATTGCAC--GAGAGGGCT
C
WT
2cpab
fad7
fad8
fad7-fad8
2cpab-fad7
2cpab-fad8
2cpab-fad7-fad8
#1
#2
#1
#2
#1
#2
kDa
25
2-Cys Prxs
Ponceau

**Supplemental Figure S1. CRISPR-based editing events of the 2-Cys Prxs genes in Arabidopsis mutant lines lacking 2-Cys Prxs and chloroplast ω-3 fatty acid desaturases (Supports Figure 1, 2 and 3). (A)** Schematic representation of *2-Cys Prx A* and *B* genes. The coding sequence and introns are indicated as black boxes and black lines, respectively. The position (red triangle) of the gRNA designed for 2-Cys PRXs genes are indicated. **(B)** Validation of homozygous mutations by Sanger sequencing. The reference sequence (*2-Cys Prx A* and *B*) and the position of the gRNAs (red line) and the protospacer adjacent motif (PAM) (blue line) are indicated. Genomic fragments containing the 2-Cys Prxs target site were obtained by PCR using oligonucleotides listed in Table S3 and sequenced. For each mutant, the obtained chromatogram is shown. Insertions and deletions are highlighted in red. **(C)** Western blot analysis of the levels of 2-Cys Prxs in wild-type and mutant lines lacking 2-Cys Prxs and/or chloroplast ω-3 fatty acid desaturases. Protein extracts (15 μg) were subjected to SDS–PAGE under reducing conditions, transferred to nitrocellulose filters and probed with anti-2-Cys Prxs antibodies. Even loading was monitored by Ponceau staining of the Rubisco large subunit. Molecular mass markers (kDa) are indicated on the right.

### Figure S2

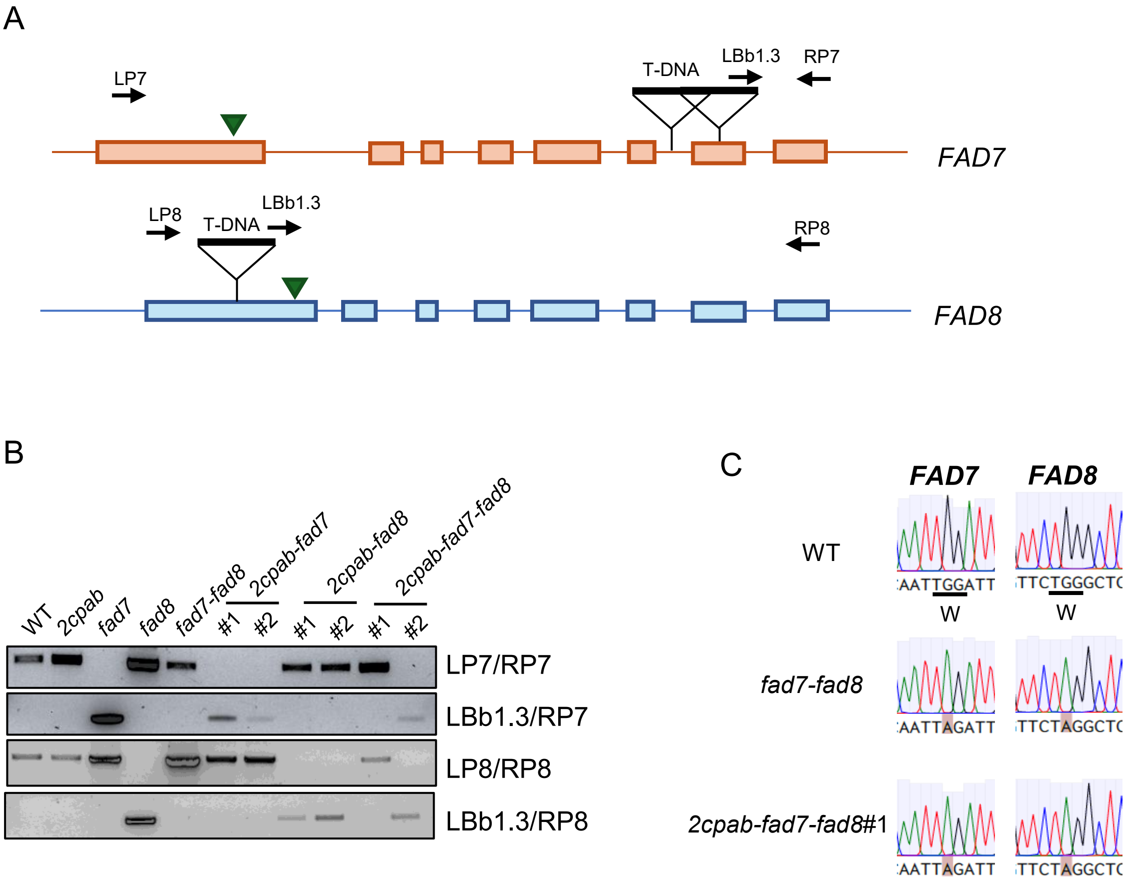
A
LP7
T-DNA
LBb1.3
RP7
FAD7
LP8
T-DNA
LBb1.3
RP8
FAD8
B
WT
2cpab
fad7
fad8
fad7-fad8
2cpab-fad7
#1
#2
2cpab-fad8
#1
#2
2cpab-fad7-fad8
#1
#2
LP7/RP7
LBb1.3/RP7
LP8/RP8
LBb1.3/RP8
C
FAD7
FAD8
WT
AATTGGATT
W
TTCTGGGCTC
W
fad7-fad8
AATTAGATT
TTCTAGGCTC
2cpab-fad7-fad8#1
AATTAGATT
TTCTAGGCTC

**Supplemental Figure S2. Genotypic characterization of chloroplast ω-3 fatty acid desaturases genes in Arabidopsis mutant lines lacking 2-Cys Prxs and chloroplast ω-3 fatty acid desaturases (Supports Figure 1, 2 and 3). (A)** Schematic representation of *FAD7* and *FAD8* genes. The coding sequence and introns are indicated as boxes and lines, respectively. The position (black inverted triangle) of the T-DNA are indicated. Green inverted triangles indicate the position of EMS mutation. The arrows indicate the primers used for genotyping, listed in Table S3. **(B)** Characterization by PCR of the T-DNA insertions. Genomic fragments containing the *FAD7* and *FAD8* genes were obtained by PCR using oligonucleotides indicated in (A) and listed in Table S3. **(C)** Validation of homozygous EMS-mutations by Sanger sequencing. The reference sequence (*FAD7* and *FAD8*) are indicated. Genomic fragments containing the *FAD7* and *FAD8* genes were obtained by PCR using oligonucleotides listed in Table S3 and sequenced. For each mutant, the obtained chromatogram is shown. Point mutations are highlighted in red.

### Figure S3

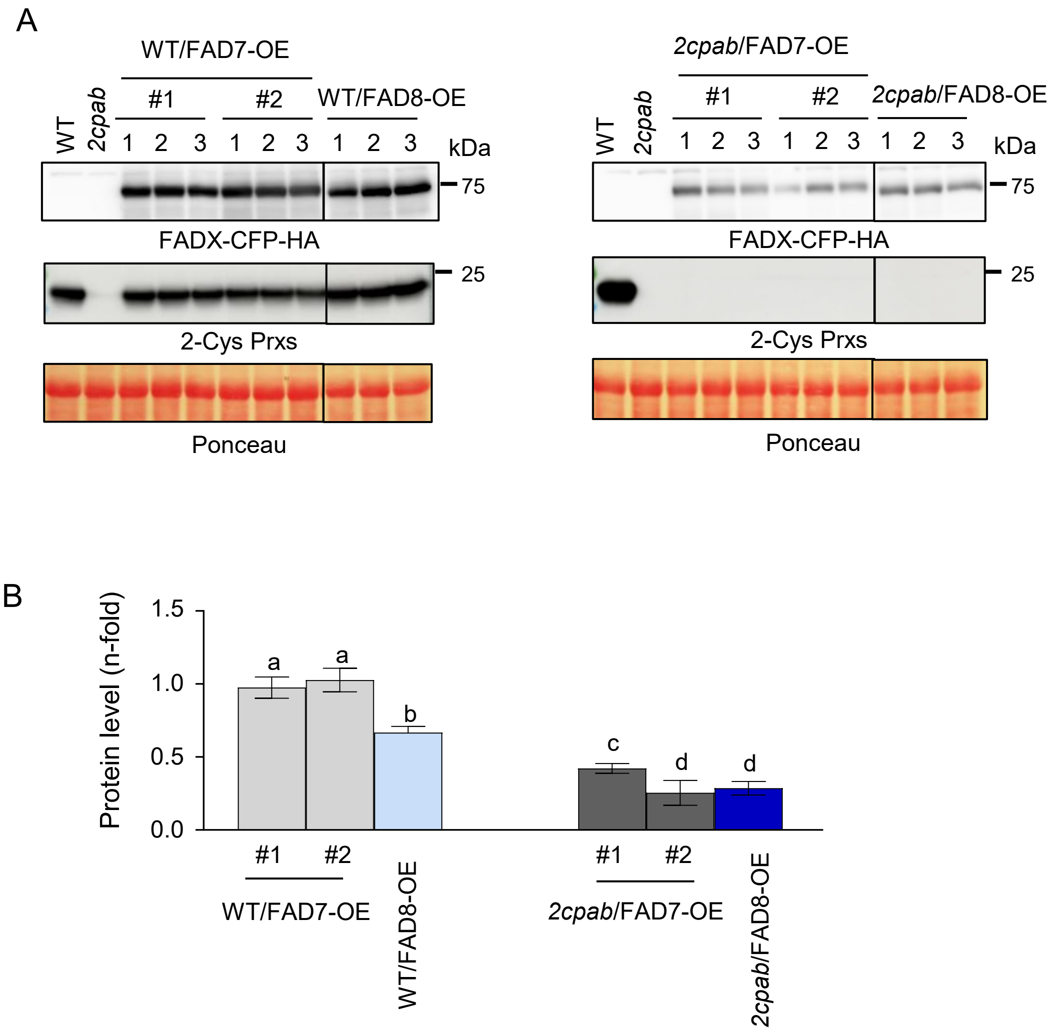
A
WT/FAD7-OE
#1
#2
WT/FAD8-OE
WT
2cpab
1 2 3 1 2 3 1 2 3
kDa
75
FADX-CFP-HA
25
2-Cys Prxs
Ponceau
2cpab/FAD7-OE
#1
#2
2cpab/FAD8-OE
WT
2cpab
1 2 3 1 2 3 1 2 3
kDa
75
FADX-CFP-HA
25
2-Cys Prxs
Ponceau
B
Protein level (n-fold)
1.5
1.0
0.5
0.0
a
a
b
c
d
d
#1
#2
WT/FAD7-OE
WT/FAD8-OE
#1
#2
2cpab/FAD7-OE
2cpab/FAD8-OE

**Supplemental Figure S3. 2-Cys Prxs protein levels in Arabidopsis leaves of transgenic lines overexpressing FAD7 and FAD8 in the WT and *2cpab* backgrounds (Supports Figure 4). (A)** Western blot analysis of the levels of CFP-HA tagged Fatty Acid Desaturase 7 (FAD7) and 8 (FAD8) and 2-Cys Prxs in protein extracts (15 µg) of rosette leaves from WT and *2cpab*, included as controls, and three individual (1, 2, and 3) from the WT/FAD7-OE#1 and #2, WT/FAD8-OE, *2cpab*/FAD7-OE#1 and #2, and *2cpab*/FAD8-CFP-HA plants, grown under LD photoperiod for 4 weeks. Even loading was monitored by Ponceau staining of the Rubisco large subunit. Molecular mass markers (kDa) are indicated on the right. **(B)** Band intensities corresponding to FAD7-CFP-HA, FAD8-CFP-HA and Rubisco large subunit in WT/FAD7-OE#1 and #2, WT/FAD8-OE, *2cpab*/FAD7-OE#1 and #2, and *2cpab*/FAD8-CFP-HA were quantified (GelAnalyzer). The contents of FAD7–CFP–HA and FAD8–CFP–HA, normalized to the levels of Rubisco large subunit, are shown relative to the levels of the WT/FAD7-CFP-HA#1 sample (arbitrarily assigned a value of 1). Data are given as the mean ± SEM of three independent replicates Letters indicate significantly different according to one-way ANOVA with a Tukey's multiple comparisons test ($p < 0.05$).

### Figure S4

A

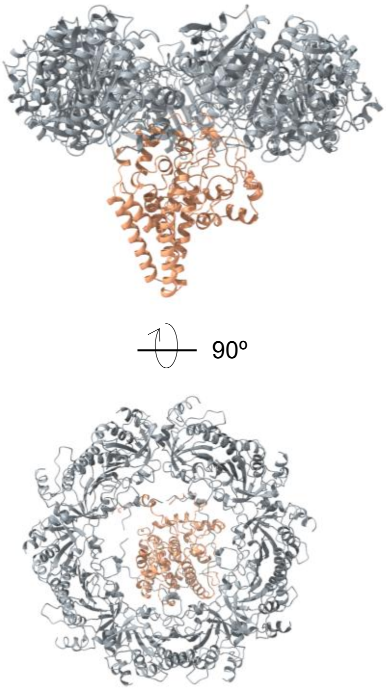


ipTM = 0.32
pTM score = 0.36

B

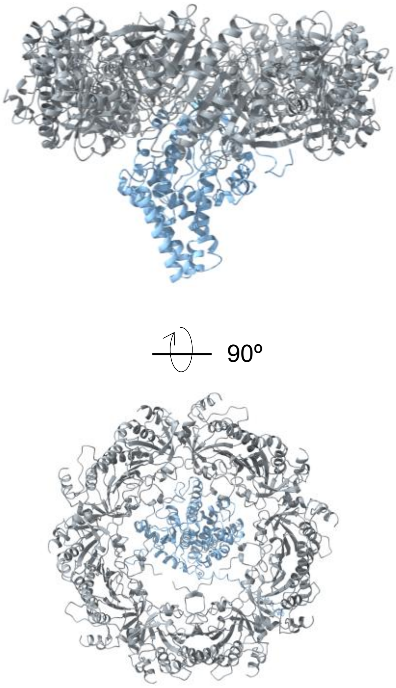


ipTM = 0.32
pTM score = 0.37

C

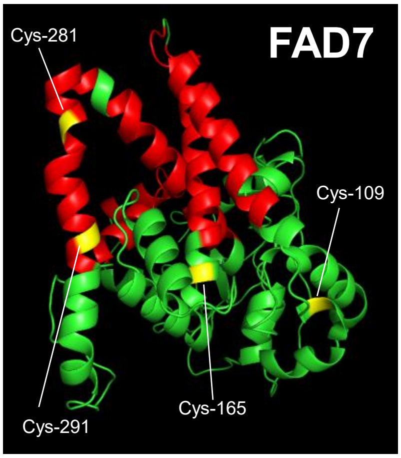


D

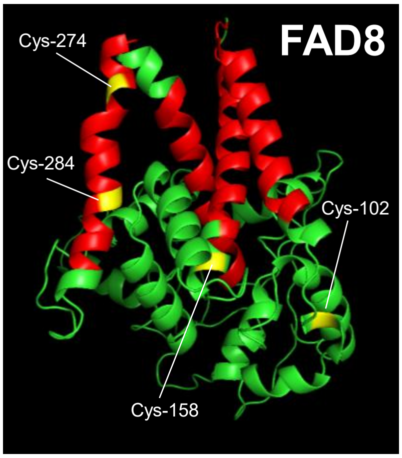

**Supplemental Figure S4. Structural modelling of chloroplast ω-3 fatty acid desaturases FAD7 and FAD8 and its interaction with 2-Cys Prxs using AlphaFold (Supports Figure 4).** Predicted structure of interaction between the 2-Cys Prx A decamer with FAD7 **(A)** and with FAD8 (**B**), using Alphafold3 prediction. The 2-Cys Prx A decamer is coloured in grey, FAD7 in orange, and FAD8 in light blue cartoons. The complexes are shown from top and side-view. Interface-predicted TM score (ipTM) and predicted Template Modeling score (pTM) are shown under the predicted structures. AlphaFold-predicted structure of FAD7 **(C)** and FAD8 **(D)**. Transmembrane domains are highlighted in red, while hydrophobic residues are shown in green. Cysteine residues are indicated and labelled within the structure. The protein structure was predicted using AlphaFold and visualized with ChimeraX.

### Figure S5

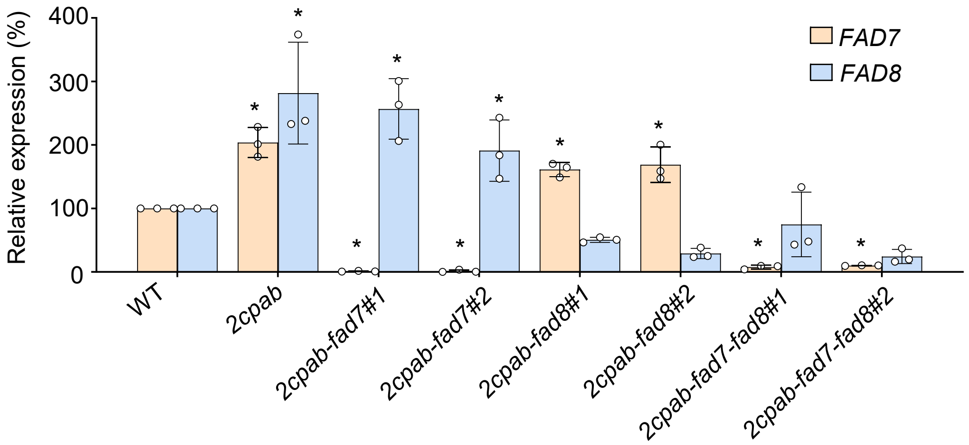


**Supplemental Figure S5. RT-qPCR expression analysis of chloroplast ω-3 desaturases in Arabidopsis wild-type and mutant lines lacking 2-Cys Prxs and chloroplast ω-3 fatty acid desaturases (Supports Figure 6 and 7).** Transcript levels of genes encoding FAD7 and FAD8 desaturases in rosette leaves from wild-type (WT) and mutant lines, as indicated, grown under long-day condition at a light intensity of 125 µE $m^{-2}$ $s^{-1}$ for 4 weeks. For each gene, the levels of transcripts were normalized against two reference genes (Material and Methods) and referenced against the levels in WT plants (arbitrarily considered as 100). Values represent the mean ± SD of three technical replicates. * Indicates significantly different to WT by one-way ANOVA with a Tukey's multiple comparison test ($p < 0.05$).

### Figure S6

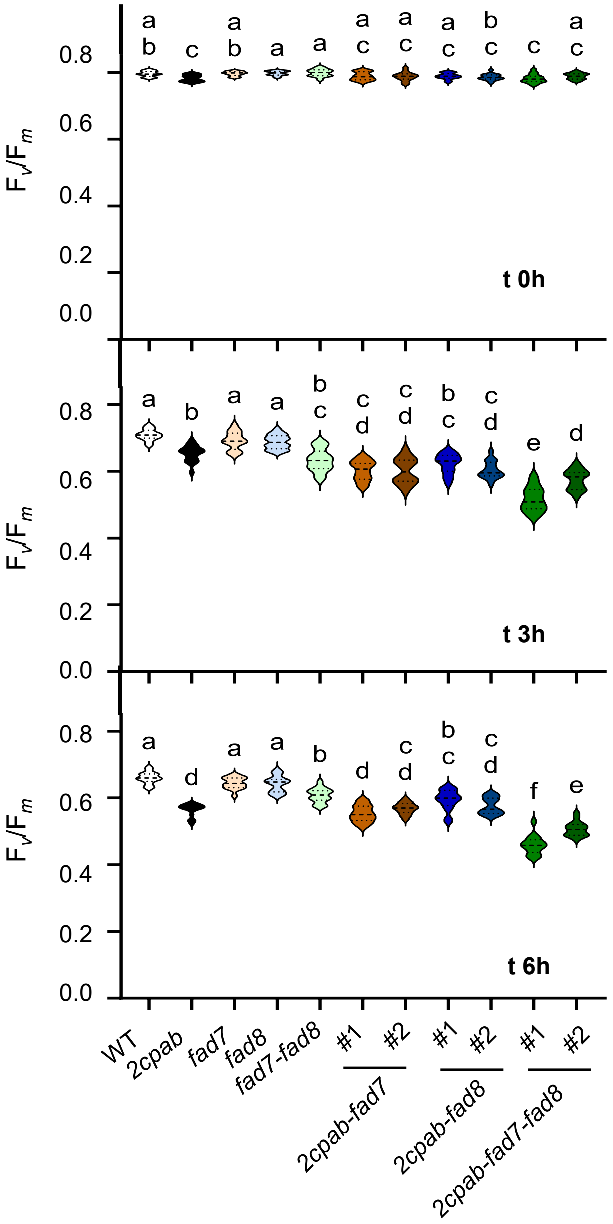


**Supplemental Figure S6. Effect of high light on photosynthetic performance in Arabidopsis leaves from WT and mutants lacking 2-Cys Prxs and/or chloroplast ω-3 fatty acid desaturases (Supports Figure 8).** Plants of the wild-type (WT) and mutant lines, as indicated, grown under long-day condition at a light intensity of 125 µE $m^{-2}$ $s^{-1}$ for 3 weeks were exposed to a light intensity of 900 µE $m^{-2}$ $s^{-1}$ at 15 ºC for 24 hours. Maximum PSII quantum yield of dark-adapted leaves represented as the ratio of variable fluorescence ($F_v$) to maximal fluorescence ($F_m$), $F_v/F_m$, was determined before (t 0h) and after 3 (t 3h) and 6 (t 6h) hours of high light treatment. Data are means ± SD of four independent plants. Letters indicate significantly different according to one-way ANOVA with a Tukey's multiple comparisons test ($p < 0.05$).
